# Modeling the cell biology of PEX11β deficiency during human neurogenesis

**DOI:** 10.64898/2026.02.13.705842

**Authors:** C Bodnya, RP Theart, V Gama

## Abstract

Peroxisomes are highly specialized organelles that are important for various metabolic functions, including β-oxidation of very-long-chain fatty acids and the synthesis of plasmalogens. Mutations in peroxisomal biogenesis proteins cause Zellweger spectrum disorders (ZSD), rare multisystem disorders often associated with neurological phenotypes. Unlike other peroxisome biogenesis proteins, PEX11β regulates peroxisomal fission, and PEX11β mutations result in milder metabolic phenotypes but persistent neurodevelopmental abnormalities, suggesting a role for PEX11β in neurodevelopment. To model PEX11β deficiency during human neurogenesis, we generated PEX11β knockout human iPSCs and differentiated them into neural progenitors and neural rosettes. PEX11β loss caused elongated peroxisomal morphology, reduced fission, and impaired recruitment of fission proteins, without affecting mitochondrial morphology or respiration. Elongated peroxisomal morphology was independent of the peroxisome-endoplasmic reticulum tether. Lipidomic analysis revealed reduced ether-linked phospholipids in PEX11β-deficient neural progenitors, suggesting impaired peroxisomal function. Finally, PEX11β deficiency led to increased neural rosette lumen size and neural progenitor number.

## Introduction

Peroxisomes are specialized organelles that are essential for lipid metabolism, including the β-oxidation of very-long-chain fatty acids (VLCFAs) and the synthesis of critical metabolites, including bile acid intermediates, cholesterol, docosahexaenoic acid, and plasmalogens (Brites et al., 2004; Ferdinandusse et al., 2009; Honsho and Fujiki, 2023; Krisans, 1992; Lazarow, 1978; Su et al., 2001). These metabolic functions are particularly important in the brain, where peroxisomal processes are key for myelin production and neural membrane integrity (Bottelbergs et al., 2010; Hulshagen et al., 2008; Kassmann, 2014; Kassmann et al., 2007; Teigler et al., 2009). Dysfunction in any of the 13 peroxisomal biogenesis proteins (PEXs) leads to defects in peroxisome membrane assembly or matrix protein import, and dysregulation of peroxisomal metabolism, resulting in a family of autosomal recessive disorders known as Zellweger spectrum disorders (ZSDs). Individuals with ZSDs present with metabolic dysfunction, such as accumulation of VLCFAs and reduction in plasmalogens, and multi-system phenotypes, including severe neurological symptoms like developmental delay and seizures (Steinberg et al., 1993). Individuals with novel variants in the peroxisomal biogenesis factor 11 beta (*PEX11β)* gene, which encodes a protein in the peroxisomal fission machinery, have been identified (Ebberink et al., 2012; Henning et al., 2024; Khoddam et al., 2024; Malekzadeh et al., 2021; Taylor et al., 2017; Tian et al., 2019; Tsinopoulou et al., 2023) (OMIM: 603867), extending the ZSD phenotype. Most reported variants in PEX11β are loss-of-function mutations. Compared to most ZSDs, PEX11β-associated disease is characterized by a milder overall phenotype and relatively subtle metabolic alterations. Yet the presence of seizures, neuropathy, and developmental delay in individuals with PEX11β mutations emphasizes the indispensable function of PEX11β during human neurodevelopment.

PEX11β is a key component of the peroxisomal fission machinery; it initiates the multi-step peroxisomal fission process in part by mediating elongation of the peroxisomal membrane via its N-terminal amphipathic helix (Delille et al., 2010; Koch et al., 2010; Schrader et al., 2012, 1998). PEX11β self-interaction has also been suggested to mediate its ability to deform the peroxisomal membrane (Bonekamp et al., 2013; Yang et al., 2025). Some of the lipids required for peroxisomal membrane expansion are proposed to originate via lipid flow from the endoplasmic reticulum (Costello et al., 2017; Hua et al., 2017), and cellular docosahexaenoic acid has also been shown to mediate peroxisomal elongation (Itoyama et al., 2012). Following peroxisomal elongation, the peroxisome undergoes constriction via mechanisms that have not been fully characterized; PEX11β may contribute to the constriction step, as it is observed at sites of peroxisomal constriction and can constrict proteo-liposomes *in vitro* (Delille et al., 2010; Schrader et al., 1998; Yoshida et al., 2015). Next, PEX11β interacts with the downstream components of the peroxisomal fission machinery: the receptors mitochondrial fission factor (MFF) and mitochondrial fission 1 (FIS1), promoting their assembly at peroxisomes (Kobayashi et al., 2007; Koch et al., 2005; Koch and Brocard, 2012). Peroxisomal fission culminates in the recruitment of dynamin-related protein 1 (DRP1), which forms oligomeric rings around the peroxisomal membrane, resulting in final membrane scission in a GTPase-dependent manner (Kamerkar et al., 2018; Koch et al., 2004, 2003). PEX11β may interact with DRP1 to enhance its GTPase activity (Williams et al., 2015). It has further been shown that PEX11β mediates peroxisomal fission independently of MFF in a FIS1- and DRP1-dependent manner (Schrader et al., 2022).

Fibroblasts isolated from individuals with PEX11β mutations display elongated peroxisomal morphology (Ebberink et al., 2012); however, the effects of PEX11β deficiency on peroxisomal morphology and function in neural cells have not been explored. Given the established link between mitochondrial morphology and neural cell fate during human neurodevelopment (Iwata et al., 2020; Khacho et al., 2016), we sought to investigate whether a similar link exists between peroxisomal morphology and neural cell fate. Mitochondrial morphology has emerged as a driver of neural cell fate decisions. In a critical post-mitotic period during neural differentiation, if mitochondria in daughter cells elongate, the cells maintain a neural progenitor identity; if mitochondria remain fragmented, the cells commit to an early-born neuronal cell fate (Iwata et al., 2020; Khacho et al., 2016). These changes in mitochondrial morphology are coupled with shifts in mitochondrial metabolism, which can also regulate the tempo of neural development (Iwata et al., 2023). However, the potential impact of peroxisomal morphology and metabolism have been overlooked in these studies. Often, perturbation of DRP1 is leveraged as a modulator of mitochondrial morphology; however, disrupting DRP1 function also impacts peroxisomal fission and potentially function. Genetic manipulation of PEX11β offers a tool to interrogate the contributions of peroxisomal morphology, without perturbing mitochondrial morphology, during neurogenesis.

The neurodevelopmental effects of depleting *Pex11β* have been examined in mouse models, where *Pex11β* deletion is lethal (Li et al., 2002). *Pex11β*-deficient mice had severe neural migration defects, despite mild metabolic changes, and died shortly after birth (Li et al., 2002). Deletion of a single allele of the *Pex11β* gene in mice resulted in differentiation delays, impaired neural migration, and neural cell death (Ahlemeyer et al., 2012). Furthermore, when *Pex11β* was knocked down in mouse embryonic stem cells, a reduction in the neural progenitor pool and impaired neural differentiation were observed after 14 days of retinoic acid treatment (Esmaeili et al., 2016). However, mouse and human neurodevelopment differ significantly, particularly in brain regions with abundant neural progenitor pools, such as the outer ventricular zone (Molnár et al., 2019). Although individuals with PEX11β mutations present with neurological symptoms, brain magnetic resonance imaging appears generally normal (Ebberink et al., 2012; Taylor et al., 2017), and the effects of PEX11β deficiency on early human neurodevelopment have not been rigorously investigated. In this study, we aimed to model the cellular biology of PEX11β deficiency in human induced pluripotent stem cell (iPSC)-derived neural cells to uncover potential links between peroxisomal morphology, peroxisomal metabolism, and neural cell fate during human neurogenesis.

## Results

### PEX11β KO iPSCs are viable and maintain stem cell identity

To model the effects of perturbing PEX11β function during neurogenesis, we knocked out *PEX11β* using CRISPR/Cas9 in human iPSCs **(Figure 1A)**. Two clones containing single-base pair deletions in the *PEX11β* gene were selected for downstream analysis after Sanger sequencing. Two isogenic unedited clones were selected and used as controls. Controls and knockouts (KOs) were confirmed using whole genome sequencing, LC-MS to detect presence vs. absence of PEX11β peptides, and RT-qPCR **(Supplementary Figures 1A-1C)**. Both control and PEX11β KO iPSCs maintained a normal karyotype **(Supplementary Figure 1D)**. Expression of the two additional PEX11 isoforms, PEX11α and PEX11γ, was unchanged in PEX11β KO iPSCs compared to control **(Supplementary Figure 1A).** Next, we validated whether knocking out PEX11β altered the stem cell identity of iPSCs. We first evaluated pluripotency by comparing the gene expression profiles of control and PEX11β KO clones with a reference data set of over 450 genome-wide transcriptional profiles of validated pluripotent cells (PluriTest). Both control and PEX11β KO clones exhibited a transcriptional profile consistent with that of a pluripotent cell **(Supplementary Figure 2A)**. To complement the pluripotency test, we evaluated levels of the key stem cell identity markers *octamer-binding transcription factor 4 (OCT4)*, *NANOG*, and *sex determining region Y-box 2 (SOX2)* using RT-qPCR, and of OCT4, NANOG, and SOX2 by immunofluorescence. There were no significant differences in stem cell identity markers between control and PEX11β KO iPSCs **(Figure 1B, Supplementary Figure 2B)**. To determine whether control and PEX11β KO iPSCs differentiated into the three germ layers, we performed trilineage differentiation. Commitment to the endoderm lineage was evaluated using SRY-box transcription factor 17 (SOX17) and forkhead box A2 (FOXA2); commitment to the mesoderm lineage was evaluated using brachyury and neural cell adhesion molecule (NCAM); and commitment to the ectoderm lineage was evaluated using paired box 6 (PAX6) and Nestin. There were no significant differences in the ability of control and PEX11β KO iPSCs to differentiate into the three germ layers **(Supplementary Figure 3)**. These findings indicate that PEX11β KO iPSCs retain stem cell identity.

**Figure 1.**
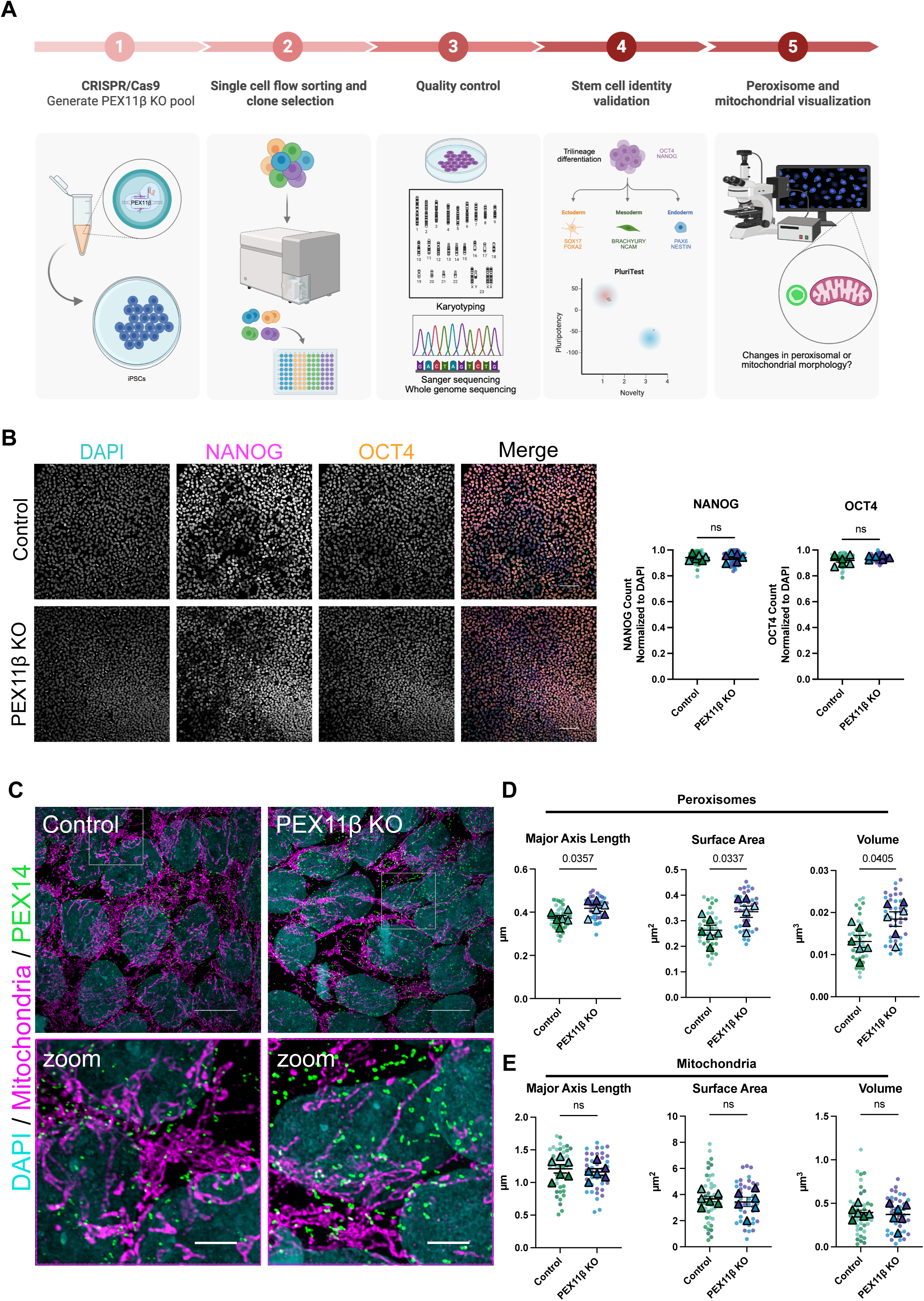
PEX11β KO iPSCs maintain stem cell identity and have elongated peroxisomal morphology. **(A)** Workflow of generating and validating PEX11β KO iPSCs. **(B)** Representative maximum intensity projections and quantification of control and PEX11β iPSCs stained for stem cell identity markers NANOG and OCT4. Scale bars = 100μm. (Spinning disk confocal [SDC]). **(C)** Representative maximum intensity projections of peroxisomal morphology (PEX14) and mitochondrial morphology (pan-mitochondria) in control and PEX11β KO iPSCs. Scale bars = 10μm. Zoom scale bars = 2.5μm (SoRa). **(D)** Quantification of peroxisomal major axis length, surface area, and volume in control and PEX11β KO iPSCs. Each dot represents the average per field of view (7 per n), each triangle represents a biological replicate, and different colors represent different clones (n=3). Analyzed using Welch’s t-test; error bars represent mean ± SEM. **(E)** Quantification of mitochondrial major axis length, surface area, and volume in control and PEX11β KO iPSCs. Each dot represents the average per field of view (7 per n), each triangle represents a biological replicate, and different colors represent different clones (n=3). Analyzed using Welch’s t-test; error bars represent mean ± SEM.

### PEX11β deficiency leads to elongated peroxisomal morphology in iPSCs, with no changes in mitochondrial morphology

PEX11β deficiency causes peroxisomal elongation in mouse embryonic fibroblasts and in fibroblasts from individuals with PEX11β mutations (Ebberink et al., 2012; Li and Gould, 2002). To test whether loss of PEX11β similarly alters peroxisome morphology in iPSCs, we stained PEX11β KO iPSCs for PEX14, a core component of the peroxisomal matrix protein import machinery (Yamashita et al., 2020). After acquiring super-resolution images, we quantified peroxisomal major axis length, surface area, and volume by briefly segmenting peroxisomes in 3D and extracting data from the resulting 3D mask (Joshi et al., 2020; Rasmussen et al., 2020; Robertson et al., 2023). PEX11β KO iPSCs displayed elongated peroxisomal morphology compared to controls, consistent with previous observations in other systems **(Figures 1C-1D, Supplementary Figure 2C)**. Because the mitochondrial and peroxisomal fission machineries and metabolic functions are tightly interconnected (Demarquoy and Le Borgne, 2015; Schrader, 2006; Schrader et al., 2015; Subramani et al., 2023), we examined whether knocking out PEX11β alters mitochondrial morphology. Mitochondrial morphology was examined in PEX11β KO iPSCs using a pan-mitochondrial antibody and super-resolution imaging, with quantification of major axis length, surface area, and volume as described above. PEX11β KO iPSCs exhibited no detectable changes in mitochondrial morphology compared to control iPSCs **(Figures 1C and 1E, Supplementary Figure 2C),** indicating that PEX11β loss selectively affects peroxisomes without altering mitochondrial morphology. This finding is essential, as a key limitation of studies that perturb both mitochondrial and peroxisomal fission proteins is that the unique effects of perturbations on peroxisomal morphology and function are often unexplored.

### PEX11β−deficient cells exhibit slight neural identity changes in early neural development

To establish whether PEX11β deficiency affects neural cell fate, we differentiated PEX11β KO iPSCs into neural progenitors (NPCs) over the course of 8 days using dual SMAD inhibition (Chambers et al., 2009) **(Figure 2A)**. PEX11β KO NPCs had no changes in the expression of the neural progenitor identity markers *PAX6, Nestin,* or *SOX2,* and the intermediate progenitor marker *t-brain gene-2* (*TBR2)* compared to control NPCs **(Figures 2B-2D).** However, the expression of *beta-3-tubulin* (*TUBB3)*, a marker of early-born neurons, was reduced in PEX11β KO NPCs compared to control NPCs **(Figure 2E)**, indicating a transient delay in neuronal differentiation. A similar phenotype is observed when mitochondrial fission is inhibited (Iwata et al., 2020; Khacho et al., 2016). However, by day 12 of NPC differentiation, there were no longer differences in expression of *TUBB3*, or other neural identity markers **(Figures 2F-2H).** Therefore, though PEX11β deficiency has an early effect on *TUBB3* expression, PEX11β-deficient NPCs recover from or compensate for this effect as they differentiate. These findings contrast with previous observations in Pex11β-deficient mice, where depletion in the neural progenitor pool and differentiation defects have been reported (Ahlemeyer et al., 2012; Esmaeili et al., 2016; Li et al., 2002), highlighting potential species-specific requirements for PEX11β during neurogenesis.

**Figure 2.**
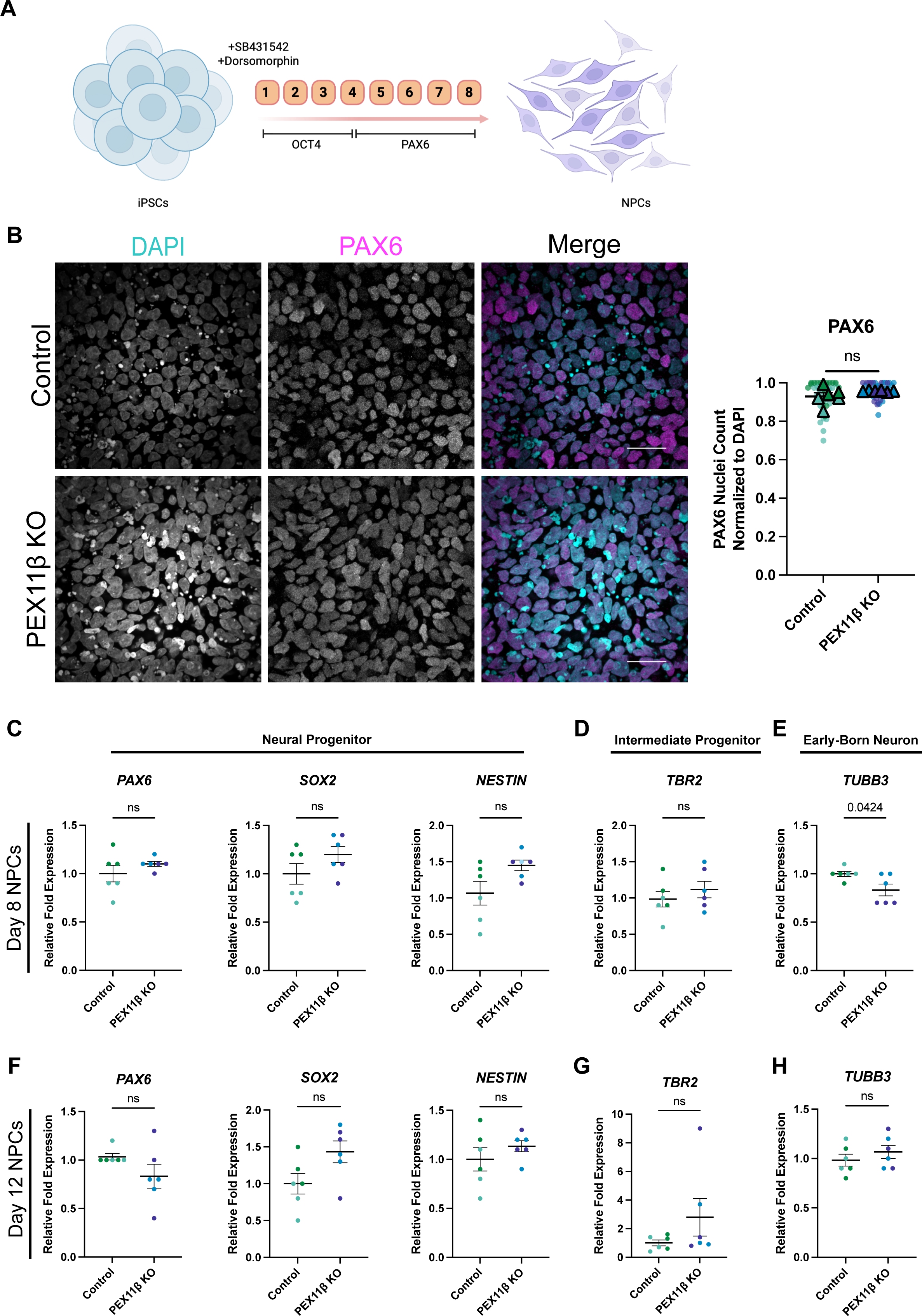
PEX11β KO iPSCs differentiate into neural progenitors, with slight changes during early neural differentiation. **(A)** Schematic of neural progenitor (NPC) differentiation using dual-SMAD inhibition. **(B)** Representative maximum intensity projections and quantification of control and PEX11β NPCs at day 8 of differentiation, stained for the neural progenitor identity marker PAX6. Scale bars = 25μm (SDC). **(C)** qRT-PCR analysis of neural progenitor identity markers *PAX6, SOX2,* and *NESTIN;* intermediate progenitor identity marker *TBR2*; and early born neuron marker *TUBB3* at day 8 of NPC differentiation. PEX11β KO NPCs were analyzed relative to control and normalized to two housekeeping genes (*GPI* and *GAPDH)*. **(D)** qRT-PCR analysis of *PAX6, SOX2,* and *NESTIN; TBR2*; and *TUBB3* at day 12 of NPC differentiation. PEX11β KO NPCs were analyzed relative to control and normalized to two housekeeping genes (*GPI* and *GAPDH)*. For qRT-PCRs, each dot represents a biological replicate, consisting of the average of two technical replicates; different colors represent different clones (n=3). Analyzed using Welch’s t-test; error bars represent mean ± SEM.

### PEX11β deficiency results in elongated peroxisomal morphology in neural progenitors

To determine whether PEX11β deficiency leads to peroxisomal elongation in iPSC-derived NPCs, we stained peroxisomes for the 70-kDa peroxisomal membrane protein, PMP70, a major component of peroxisomal membranes in mature, import-competent peroxisomes (Imanaka et al., 2000); PEX14 (Yamashita et al., 2020); and catalase, a key enzyme that breaks down the hydrogen peroxide formed as a byproduct of the first step of peroxisomal β-oxidation (Hashimoto and Hayashi, 1987). PEX11β KO NPCs had elongated peroxisomal morphology compared to control NPCs when visualized with both peroxisomal membrane and peroxisomal matrix markers **(Figures 3A-3F)**, demonstrating that PEX11β deficiency results in peroxisomal elongation that is maintained with neural differentiation.

**Figure 3.**
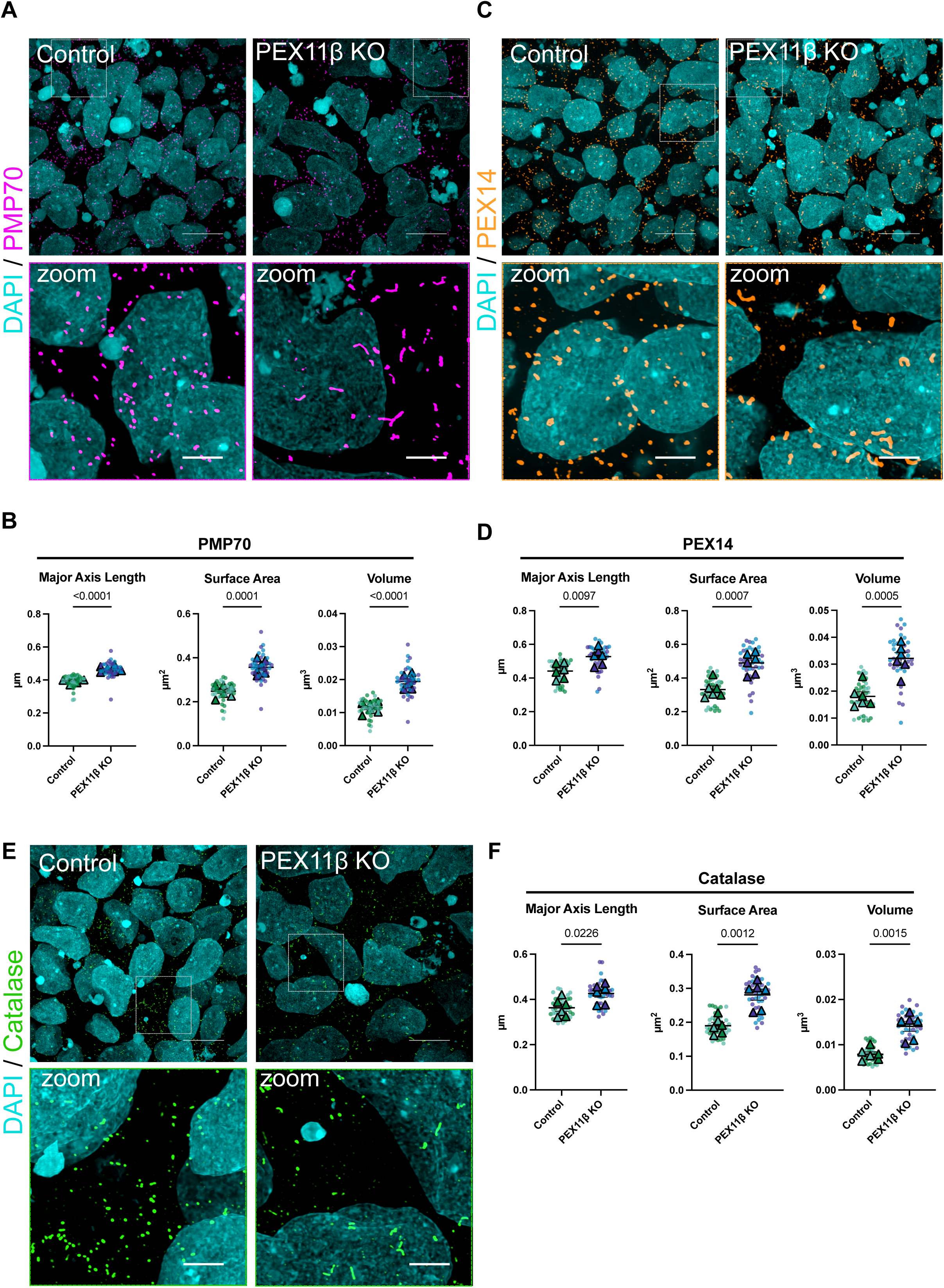
PEX11β KO NPCs have elongated peroxisomal morphology. **(A)** Representative maximum intensity projections of peroxisomal morphology (PMP70) in control and PEX11β KO NPCs at day 8 of differentiation. Scale bars = 10μm. Zoom scale bars = 2.5μm (SoRa). **(B)** Quantification of peroxisomal major axis length, surface area, and volume in control and PEX11β KO NPCs (PMP70). **(C)** Representative maximum intensity projections of peroxisomal morphology (PEX14) in control and PEX11β KO NPCs at day 8 of differentiation. Scale bars = 10μm. Zoom scale bars = 2.5μm (SoRa). **(D)** Quantification of peroxisomal major axis length, surface area, and volume in control and PEX11β KO NPCs (PEX14). **(E)** Representative maximum intensity projections of peroxisomal morphology (catalase) in control and PEX11β KO NPCs at day 8 of differentiation. Scale bars = 10μm. Zoom scale bars = 2.5μm (SoRa). **(F)** Quantification of peroxisomal major axis length, surface area, and volume in control and PEX11β KO NPCs (catalase). For all graphs, each dot represents the average per field of view (7 per n), each triangle represents a biological replicate, and different colors represent different clones (n=3). Analyzed using Welch’s t-test; error bars represent mean ± SEM.

### Changes in peroxisomal number and morphology are differentiation dependent

Because peroxisomal fission is a mechanism by which cells maintain peroxisome number, cells deficient in PEX11β have been previously reported to have a reduced number of peroxisomes, in addition to their elongated morphology (Ahlemeyer et al., 2012; Ebberink et al., 2012; Yang et al., 2025). Notably, PEX11β KO iPSCs do not have fewer peroxisomes than control iPSCs **(Supplementary Figures 4A, 4B)**, while PEX11β KO NPCs have significantly fewer peroxisomes than control NPCs **(Supplementary Figures 4C, 4D)**. To our knowledge, this is one of the first reports of a PEX11β-deficient model that shows elongated peroxisomal morphology without a significant reduction in peroxisome number. However, this effect was differentiation-dependent, as peroxisome number was reduced in PEX11β KO NPCs but not in iPSCs. Therefore, we investigated how peroxisomal morphology, number, and *PEX11β* mRNA expression change in control cells during an 8-day differentiation from iPSCs into NPCs. Peroxisome major axis length, surface area, and volume were significantly increased at day 6 of differentiation and declined by day 8 (**Figures 4A, 4B)**. Peroxisome number decreased by nearly half within the first two days of NPC differentiation and remained significantly lower than in iPSCs throughout the differentiation period **(Figure 4C)**. These dynamic changes in peroxisomal morphology coincided with peaks in *PEX11β* mRNA expression at days 4 and 6 of differentiation **(Figure 4D)**, and with maximal expression of the early neural progenitor marker PAX6 at days 6 and 8 of differentiation **(Figure 4E)**. Together, these data show that peroxisomal remodeling and *PEX11β* expression are tightly linked to acquisition of neural progenitor identity, suggesting a requirement for PEX11β in peroxisomal fission and proliferation in neural cells but not in iPSCs.

**Figure 4.**
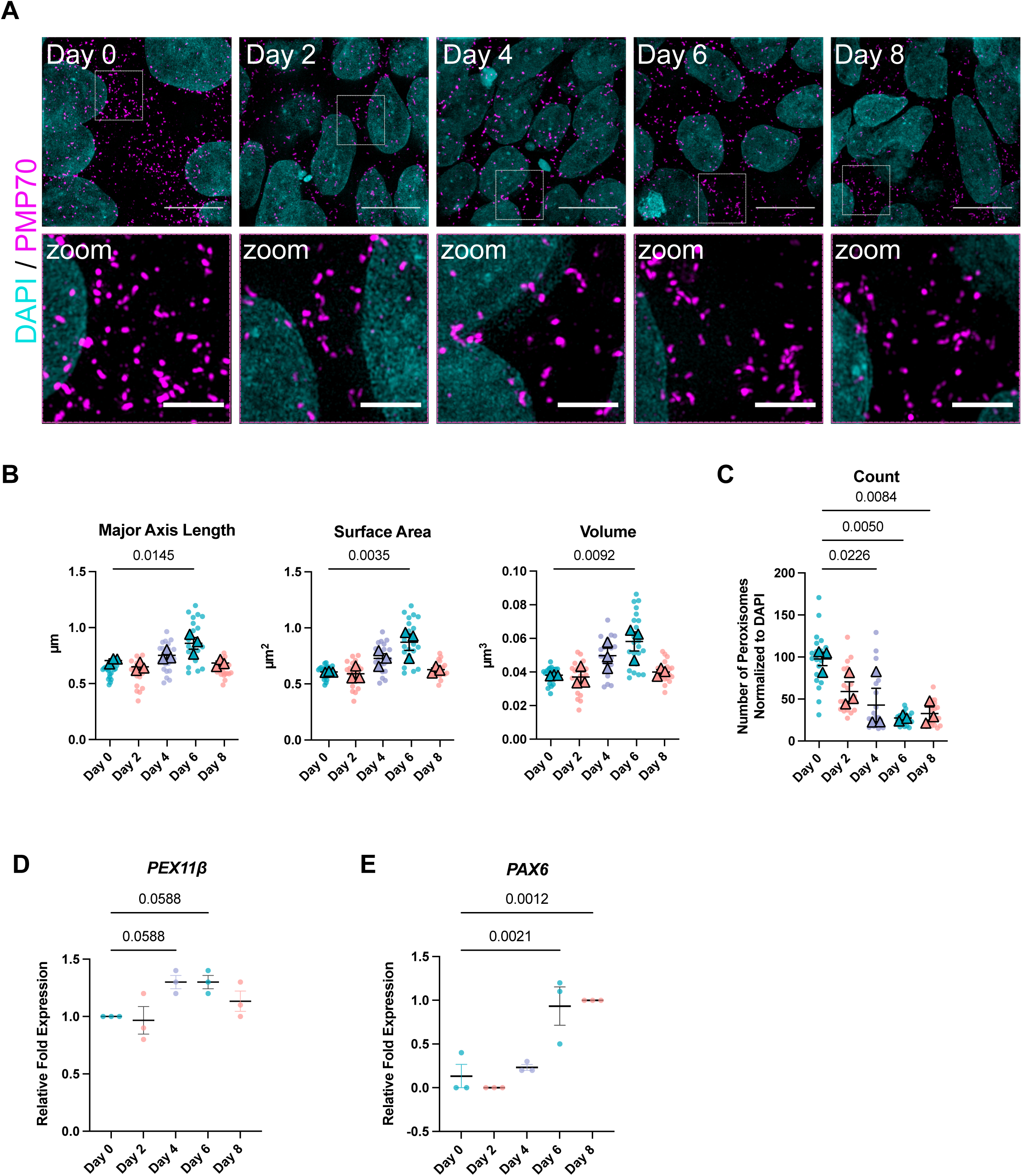
Peroxisome morphology and number change during NPC differentiation. **(A)** Representative maximum intensity projections of peroxisomal morphology (PMP70) in control NPCs imaged every two days of differentiation for 8 days. Scale bars = 10μm. Zoom scale bars = 2.5μm (SIM). **(B)** Quantification of peroxisomal major axis length, surface area, and volume in control NPCs over 8 days of differentiation. Each dot represents average per field of view (7 per n); each triangle represents the mean of the biological replicate (n=3). Analyzed using one-way ANOVA with Dunnett’s multiple comparisons; error bars represent mean ± SEM. **(C)** Quantification of peroxisomal count normalized to DAPI in control NPCs over 8 days of differentiation. Each dot represents average per field of view (7 per n); each triangle represents the mean of the biological replicate (n=3). Analyzed using one-way ANOVA with Dunnett’s multiple comparisons; error bars represent mean ± SEM. **(D)** qRT-PCR analysis of *PEX11β* gene expression during 8-day NPC differentiation. Analyzed relative to day 0 and normalized to two housekeeping genes (*GPI* and *GAPDH)*. Each dot on the graph represents a biological replicate. Analyzed using one-way ANOVA with Dunnett’s multiple comparisons; error bars represent mean ± SEM. **(E)** qRT-PCR analysis of *PAX6* gene expression during 8-day NPC differentiation, relative to day 8 of differentiation and normalized to two housekeeping genes (*GPI* and *GAPDH)*. Each dot on the graph represents a biological replicate. Analyzed using one-way ANOVA with Dunnett’s multiple comparisons; error bars represent mean ± SEM.

### The downstream peroxisomal fission machinery is perturbed in PEX11β KO neural progenitors

Given our observation that NPCs may be more reliant on PEX11β-mediated fission than iPSCs, we tested whether PEX11β deficiency reduced the frequency of peroxisomal fission events in NPCs. To examine this, we live-imaged NPCs after transduction with CellLight Peroxisome-GFP, a BacMam vector that encodes GFP linked to the peroxisomal C-terminal targeting sequence (PTS1). We then analyzed the frequency of peroxisomal fission events in control and PEX11β KO NPCs using the Mitochondrial Event Localizer (MEL) algorithm (Theart et al., 2020), applied here as a peroxisomal event localizer (PEL), MEL/PEL processes time-lapse 3D z-stack images by operating on 3D instance-labeled organelle masks to detect topological split and merge events. The algorithm outputs per-frame fission event counts together with 3D event locations indicating where peroxisomal fission events are most likely to have occurred at distinct time points. Importantly, MEL/PEL is organelle-agnostic and does not rely on mitochondria-specific assumptions. Briefly, a fission event was defined when a single labeled peroxisomal structure at time *t* split into two (or more) distinct daughter structures at *t+1* and remained split at *t+2* (a conservative persistence criterion to suppress transient segmentation artifacts). Using this approach, live-cell imaging and PEL quantification of control and PEX11β KO NPCs showed that peroxisomal fission events were significantly reduced in PEX11β KO NPCs compared to control NPCs **(Supplementary Figures 5A-5C, Supplementary Videos 1, 2)**. To identify whether PEX11β deficiency alters the recruitment of the downstream components of the peroxisomal fission machinery to peroxisomes, we imaged NPCs at day 8 of differentiation and quantified colocalization between PMP70 and MFF positive (+) or FIS1+ puncta. Because PEX11β-deficient NPCs also have a significantly reduced number of peroxisomes, quantification was normalized to the total number of MFF+ or FIS1+ puncta, thereby showing the proportion of total MFF or FIS1 that colocalized with PMP70. PEX11β KO NPCs displayed reduced colocalization between MFF and PMP70, as well as between FIS1 and PMP70, when compared to control NPCs **(Figures 5A, 5B, 5D, 5E, Supplementary Figures 6A-6B).** Because colocalization between PMP70 and MFF and FIS1, the receptors for DRP1 was decreased, we next tested whether there were changes in colocalization between PMP70 and DRP1+ puncta. PEX11β KO NPCs showed reduced colocalization between PMP70 and DRP1 **(Figures 5G, 5H, Supplementary Figure 6C)**. These findings support prior work identifying PEX11β as a key factor in recruiting MFF and FIS1 (Kobayashi et al., 2007; Koch and Brocard, 2012; Schrader et al., 2016, 2012). Given that MFF, FIS1, and DRP1 are shared components of the mitochondrial and peroxisomal fission systems (Schrader, 2006; Subramani et al., 2023), colocalization of these proteins with mitochondria was also assessed. No significant changes in the association of MFF, FIS1, or DRP1 with mitochondria were detected **(Figures 5C, 5F, 5I)**, indicating that the basal mitochondrial fission machinery remains intact in PEX11β-deficient cells. These results align with the absence of changes in mitochondrial network morphology in PEX11β KO iPSCs.

**Figure 5.**
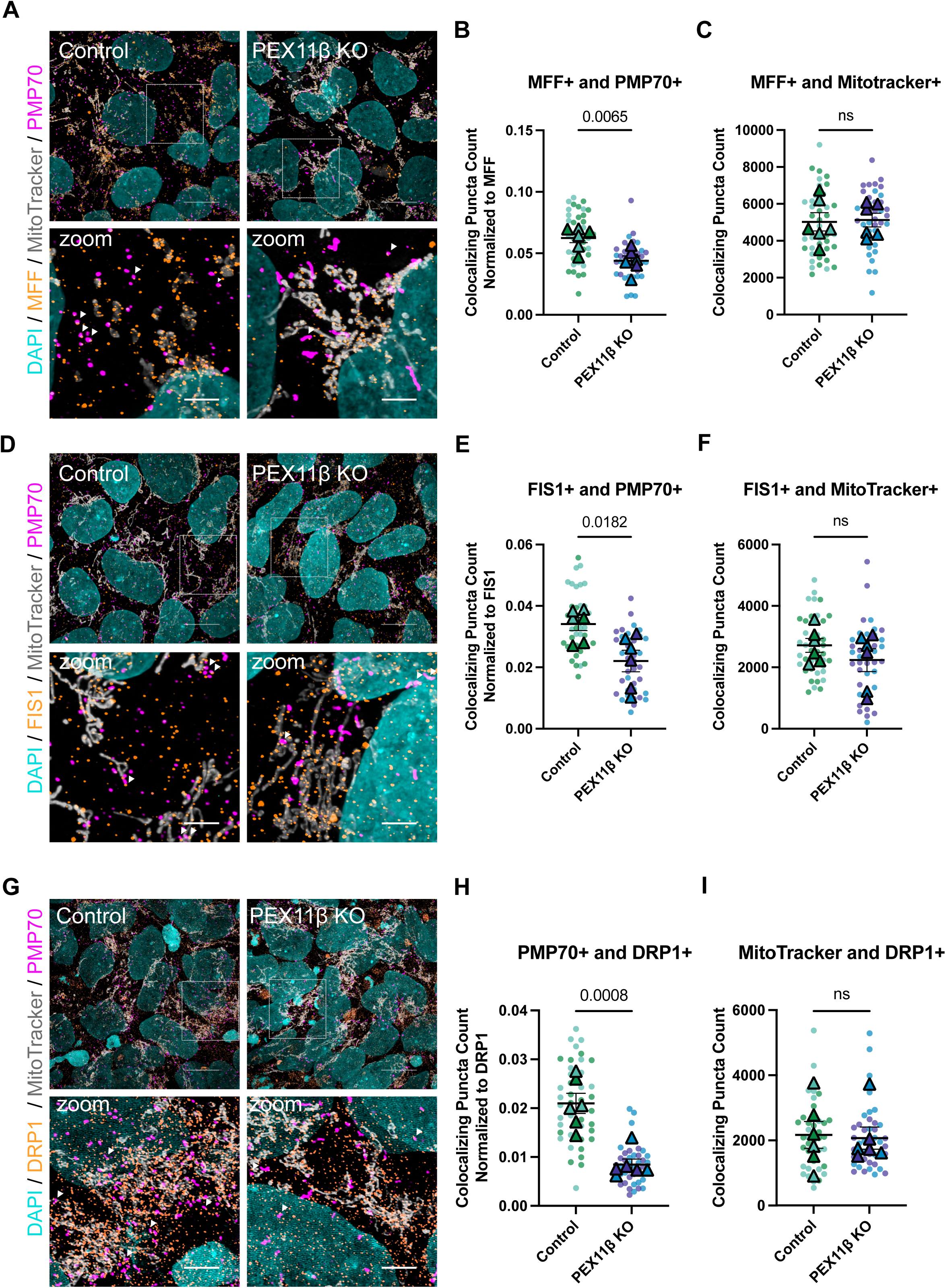
Colocalization with downstream peroxisomal fission factors is reduced in PEX11β KO NPCs. **A)** Representative maximum intensity projections of MFF, mitochondria (MitoTracker), and peroxisomes (PMP70) in control and PEX11β NPCs at day 8 of differentiation. Scale bars = 10μm. Zoom scale bars = 2.5μm (SoRa). **(B)** Quantification of MFF+ puncta that colocalize with PMP70+ puncta, normalized to total number of MFF+ puncta. **(C)** Quantification of MFF+ puncta that overlap with MitoTracker. **(D)** Representative maximum intensity projections of FIS1, mitochondria (MitoTracker), and peroxisomes (PMP70) in control and PEX11β NPCs at day 8 of differentiation. Scale bars = 10μm. Zoom scale bars = 2.5μm (SoRa). **(E)** Quantification of FIS1+ puncta that colocalize with PMP70+ puncta, normalized to total number of FIS1+ puncta **(F)** Quantification of FIS1+ puncta that overlap with MitoTracker**. (G)** Representative maximum intensity projections of DRP1, mitochondria (MitoTracker), and peroxisomes (PMP70) in control and PEX11β NPCs at day 8 of differentiation. Scale bars = 10μm. Zoom scale bars = 2.5μm (SoRa). **(H)** Quantification of DRP1+ puncta that colocalize with PMP70+ puncta, normalized to total number of DRP1+ puncta. **(I)** Quantification of DRP1+ puncta that overlap with MitoTracker. For all graphs, each dot represents the average per field of view (7 per n), each triangle represents a biological replicate, and different colors represent different clones (n=3). Analyzed using Welch’s t-test; error bars represent mean ± SEM.

### PEX11β deficiency does not affect mitochondrial morphology or respiration in neural progenitors

Although the mitochondrial association of fission proteins was unchanged in PEX11β-deficient NPCs, the tight metabolic connection between peroxisomes and mitochondria prompted examination of mitochondrial morphology. Mitochondria were labeled with a pan-mitochondrial marker at day 8 of differentiation, imaged by super-resolution microscopy, and quantified for major axis length, surface area, and volume. No significant differences in mitochondrial morphology were detected between PEX11β KO and control NPCs **(Figures 6A, 6B)**. To assess whether PEX11β deficiency affects mitochondrial respiration, oxygen consumption rate (OCR) and extracellular acidification rate (ECAR) were measured in day 8 NPCs using the Seahorse XF Analyzer. ECAR served as a proxy for glycolysis, whereas OCR reported oxidative phosphorylation. No differences in overall OCR or ECAR were detected between PEX11β KO and control NPCs (**Figure 6C)**. Basal, maximal, and ATP-linked respiration were likewise unchanged in PEX11β KO NPCs, indicating preserved ATP production capacity **(Figures 6D-6G)**. PEX11β KO similarly did not have changes in proton leak and coupling efficiency **(Figures 6H, 6I)**, demonstrating effective coupling of substrate oxidation to ATP synthesis. Thus, PEX11β-deficient NPCs exhibit normal mitochondrial morphology and respiration. These findings contrast with reports of secondary mitochondrial defects in Pex11β-deficient mice (Colasante et al., 2024) and mitochondrial fragmentation in PEX11β-deficient HEK293T cells carrying a pathogenic polyQ expansion (Kumar et al., 2025).

**Figure 6.**
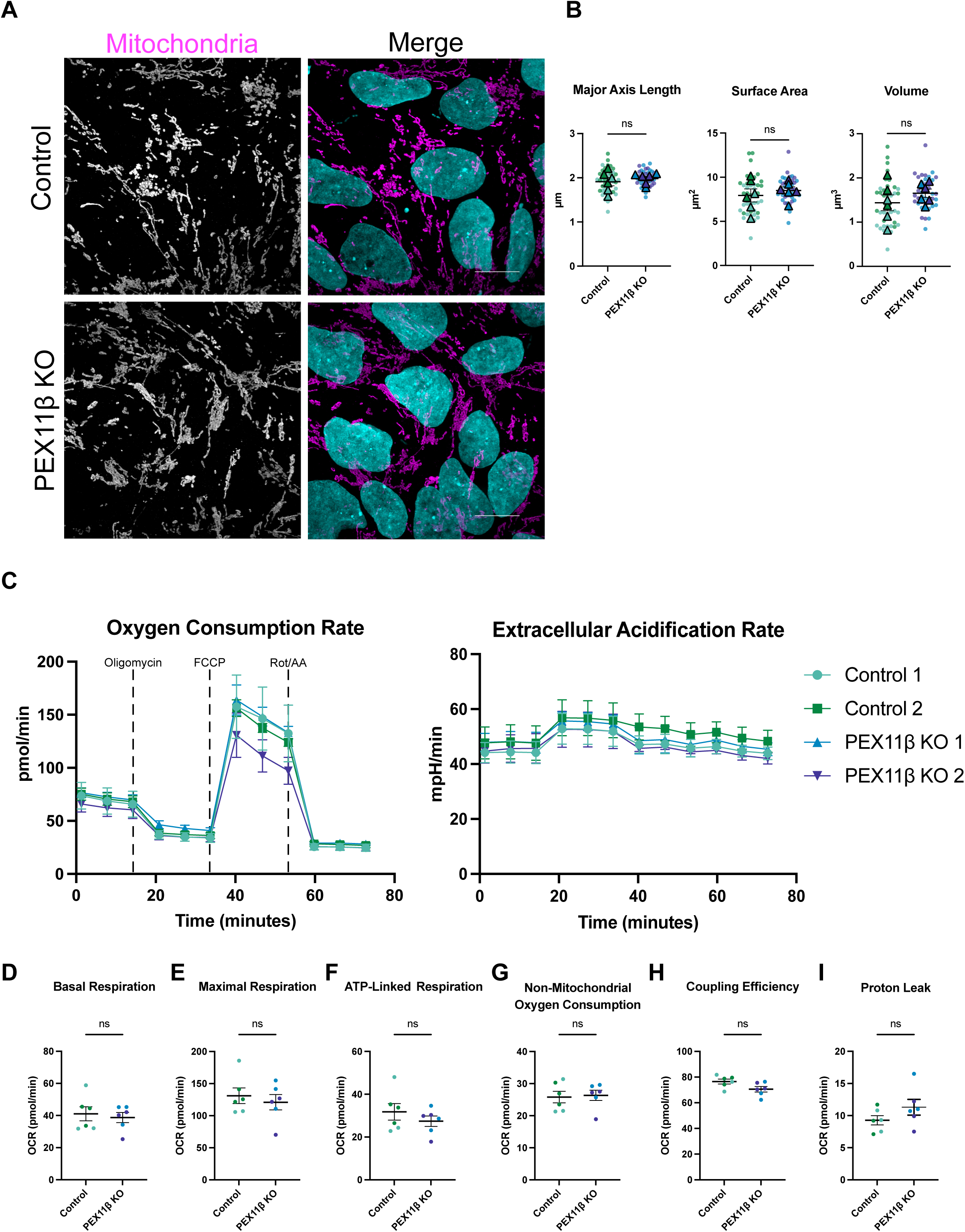
PEX11β deficiency does not impact mitochondrial morphology or mitochondrial respiration in NPCs. **A)** Representative maximum intensity projections of mitochondrial morphology in control and PEX11β KO NPCs at day 8 of differentiation. Scale bars = 10μm (SoRa). **(B)** Quantification of mitochondrial morphology in control and PEX11β KO NPCs. Each dot represents the average per field of view (7 per n), each triangle represents a biological replicate, and different colors represent different clones (n=3). Analyzed using Welch’s t-test; error bars represent mean ± SEM. **(C)** Oxygen consumption rate (OCR) and extracellular acidification rate (ECAR) in control and PEX11β KO NPCs at day 8 of differentiation (n=3 with 16 wells per replicate). Oligomycin was added at 20 minutes, FCCP was added at 40 minutes, and rotenone and antimycin A (Rot/AA) were added at 60 minutes (indicated on graph). Analyzed using two-way ANOVA followed by Šídák multiple comparisons test; error bars represent mean ± SEM. **(D)** Basal respiration in control and PEX11β KO NPCs (last rate measurement before oligomycin injection – non-mitochondrial respiration after Rot/AA injection). **(E)** Maximal respiration in control and PEX11β KO NPCs (maximal rate measurement after FCCP injection – non-mitochondrial respiration after Rot/AA injection). **(F)** ATP-linked respiration in control and PEX11β KO NPCs (last rate measurement before oligomycin injection – minimum rate measurement after oligomycin injection). **(G)** Non-mitochondrial oxygen consumption (minimum rate measurement after Rot/AA injection)**. (H)** Proton leak (minimum rate measurement after oligomycin injection – non-mitochondrial respiration after Rot/AA injection). **(I)** Coupling efficiency (ATP production rate / basal respiration rate) calculated using OCR in control and PEX11β KO NPCs. All analyzed using Welch’s t-test; error bars represent mean ± SEM.

### Perturbing the peroxisome-endoplasmic reticulum tether does not change peroxisomal morphology in PEX11β-deficient neural progenitors

Our examination of the downstream peroxisomal fission machinery revealed impaired fission and elongated peroxisomal morphology in PEX11β-deficient NPCs. Although PEX11β has been proposed to mediate both membrane elongation and division, our findings, together with prior reports, indicate that loss of PEX11β predominantly results in peroxisomal elongation. Previously, it has been suggested that this phenotype may be due to continual lipid flow from the endoplasmic reticulum (ER) to peroxisomes, which is made possible via a tether between vesicle associated membrane protein-associated protein B (VAPB) at the ER and acyl-CoA binding domain-contacting protein 5 (ACBD5) at the peroxisomes (Costello et al., 2017; Hua et al., 2017) **(Figure 7A)**. To interrogate whether peroxisome-ER contacts, and consequently, ER lipid flow, contribute to the elongated peroxisomal morphology in PEX11β-deficient cells, we disrupted this tether via RNAi-mediated knockdown of *ACBD5* in control and PEX11β KO NPCs at day 4 of differentiation. We hypothesized that if elongated peroxisomal morphology in PEX11β-deficient NPCs is a consequence of lipid flow from the ER, then *ACBD5* knockdown would result in shorter (reduced major axis length) and smaller (reduced surface area and volume) peroxisomes. After confirming efficient *ACBD5* knockdown **(Figure 7B),** peroxisomal morphology was assessed by super-resolution microscopy in regions with markedly reduced ACBD5 signal. In PEX11β KO NPCs, peroxisomal major axis length was unchanged between scramble controls and ACBD5-depleted cells **(Figures 7C, 7D)**. Notably, we found increased peroxisomal surface area and volume in PEX11β KO NPCs in which *ACBD5* was knocked down **(Figure 7D)**. Thus, elongated peroxisomal morphology in PEX11β KO NPCs does not result from ER lipid flow and contrasts with phenotypes described in other models of impaired peroxisomal morphology, including MFF- and DRP1-deficient cells (Costello et al., 2017; Darwisch et al., 2020; Hua et al., 2017). Furthermore, in control NPCs, we observed elongated peroxisomal morphology upon *ACBD5* knockdown **(Figures 7C, 7D)**, which was opposite to effects reported in non-neural systems (Costello et al., 2017; Darwisch et al., 2020; Hua et al., 2017). These findings indicate that PEX11β-dependent peroxisomal elongation in NPCs is largely independent of ER lipid transfer, while the effects of *ACBD5* knockdown on control NPCs may reflect metabolic requirements unique to neural cell types.

**Figure 7.**
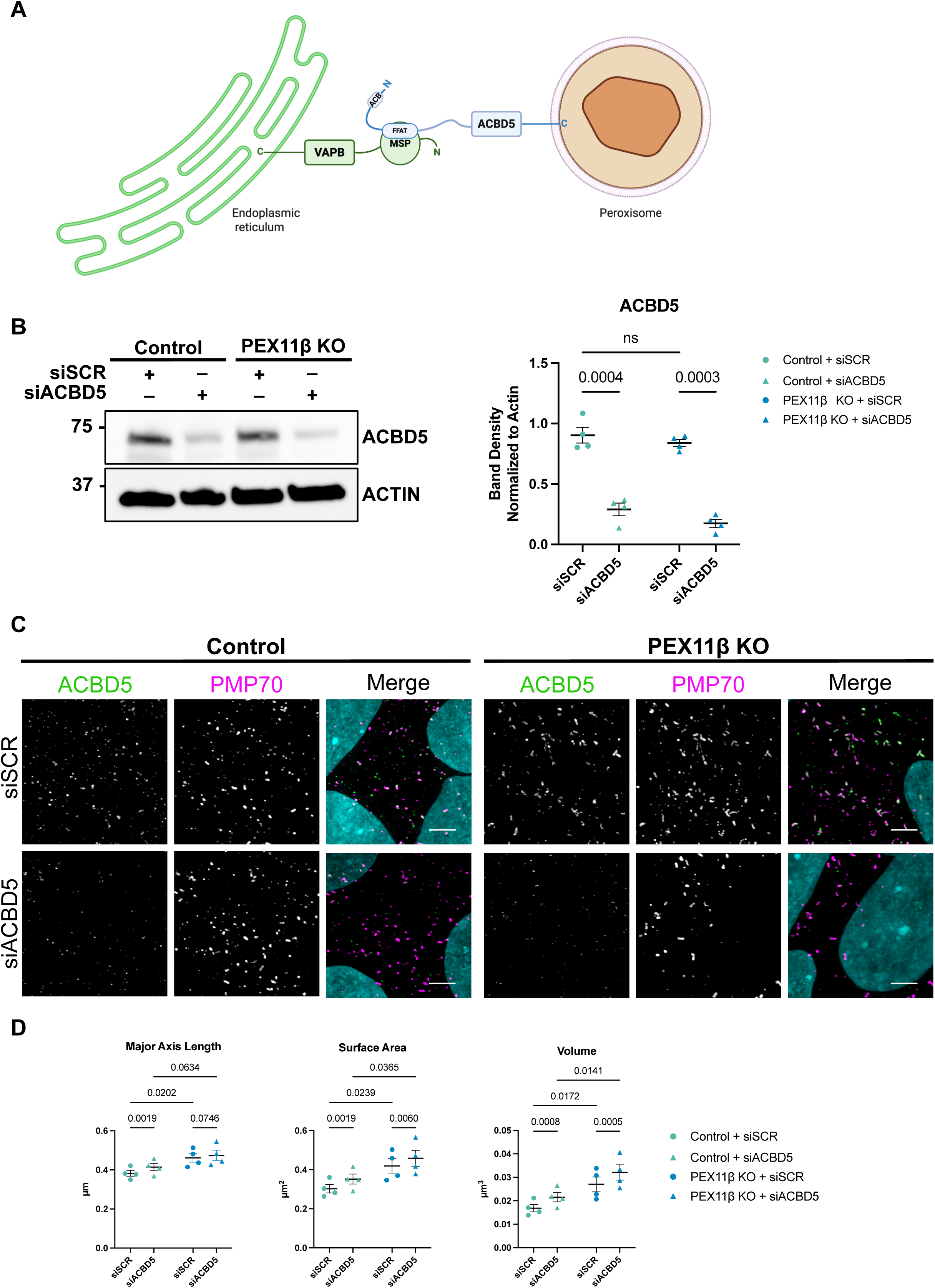
*ACBD5* knockdown does not change peroxisomal morphology in PEX11β KO NPCs. **(A)** Schematic of the interaction between ACBD5 and VAPB at peroxisome-ER contact sites, which is mediated by the FFAT motif of ACBD5 and the MSP domain of VAPB. Adapted from (Kors et al., 2022). **(B)** Representative western blot of ACBD5 knockdown in control and PEX11β KO NPCs at day 4 of differentiation and quantification of western blot band density of ACBD5 normalized to β-actin loading control. Each dot represents a biological replicate (n=4); circles represent siSCR and triangles represent siACBD5. Analyzed using two-way ANOVA with uncorrected Fisher’s LSD multiple comparisons; error bars represent mean ± SEM. **(C)** Representative maximum intensity projections of ACBD5 and PMP70 in control and PEX11β KO NPCs treated with either scramble control or *ACBD5* siRNA at day 4 of NPC differentiation. Zoom scale bars = 2.5μm (SoRa). **(D)** Quantification of peroxisomal major axis length, surface area, and volume in control and PEX11β KO NPCs at day 4 of differentiation, treated with either scramble control or *ACBD5* siRNA. Each dot represents a biological replicate (n=4); circles represent siSCR and triangles represent siACBD5. Analyzed using two-way ANOVA with uncorrected Fisher’s LSD multiple comparisons; error bars represent mean ± SEM.

### Ether-linked phospholipids are reduced in PEX11β-deficient NPCs

To assess the impact of PEX11β deficiency on lipid metabolism, global untargeted lipidomics of day 8 NPCs was performed. PEX11β KO NPCs had reduced levels of ether-linked phosphatidylcholine and phosphatidylserine species **(Table 1A, 1B)**. These results extend previous findings in PEX11β-deficient human fibroblasts, where reduced levels of phosphatidylethanolamine plasmalogens were observed (Abe et al., 2023). These findings suggest disrupted plasmalogen synthesis in PEX11β-deficient NPCs, potentially indicating impaired peroxisomal lipid metabolism (Brites et al., 2004; Honsho and Fujiki, 2023). To gain a more robust understanding of what disturbances occur at the level of the lipidome when peroxisomal fission is disturbed, we contrasted our findings in PEX11β-deficient NPCs with lipidomics data from DRP1-deficient NPCs, which have defective peroxisomal and mitochondrial fission (Baum et al., 2024; Robertson et al., 2023). Phosphatidylethanolamine and phosphatidylcholine ether-linked phospholipids were significantly reduced **(Table 2A)**, while triglyceride species with both long and very-long-chain fatty acid side chains were significantly increased in DRP1-deficient NPCs **(Table 2B)**. Both PEX11β and DRP1-deficient NPCs had an increase in sphingomyelin species, potentially containing a very-long-chain side chain **(Table 1A**, **Table 2A, 2B)**, implicating peroxisomal fission as a key determinant of sphingomyelin metabolism.

**Table 1.** PEX11β-deficient NPCs show reduced ether-linked phospholipids. (A) Fold change and p-values of significantly changed lipids between control and PEX11β KO NPCs at day 8 of differentiation in positive ion mode. Decreased lipids in blue, increased lipids in red. **(B)** Fold change and p-values of significantly changed lipids between control and PEX11β KO NPCs at day 8 of differentiation in negative ion mode. Decreased lipids in blue, increased lipids in red.

| Lipid Species | FC | log2(FC) | p-value | -log10(p) |
| --- | --- | --- | --- | --- |
| PC O-40:1 | 0.30626 | -1.7072 | 0.00046464 | 3.3329 |
| PC O-42:2 | 0.36038 | -1.4724 | 0.0018527 | 2.7322 |
| PC O-36:0 | 0.36907 | -1.438 | 0.0067067 | 2.1735 |
| PC O-34:0 | 0.37991 | -1.3963 | 0.0047526 | 2.3231 |
| LPC 16:1/0:0 | 0.39145 | -1.3531 | 0.04798 | 1.3189 |
| PC O-40:2 | 0.39755 | -1.3308 | 0.0050966 | 2.2927 |
| PC 46:3 | 0.41217 | -1.2787 | 0.0126 | 1.8996 |
| PC O-38:1 | 0.41363 | -1.2736 | 0.0079693 | 2.0986 |
| PC 25:0 | 0.42235 | -1.2435 | 0.031207 | 1.5057 |
| LPC O-18:1 | 0.43895 | -1.1879 | 0.010778 | 1.9674 |
| PC O-30:1 | 0.48317 | -1.0494 | 0.0018165 | 2.7408 |
| PC O-36:1 | 0.48371 | -1.0478 | 0.010071 | 1.9969 |
| LPE O-18:1 | 0.48826 | -1.0343 | 0.018717 | 1.7278 |
| LPC O-16:1 | 0.49944 | -1.0016 | 0.0096856 | 2.0139 |
| PC O-44:4 | 0.49975 | -1.0007 | 0.0038006 | 2.4201 |
| SM 44:5;2O\|SM 18:1;2O/26:4 | 3.7336 | 1.9006 | 0.034819 | 1.4582 |

B PEX11β KO vs Control – Negative Ion Mode
| Lipid Species | FC | log2(FC) | p-value | -log10(p) |
| --- | --- | --- | --- | --- |
| PC O-40:1\|PC O-24:1_16:0 | 0.27491 | -1.863 | 0.00058787 | 3.2307 |
| PC O-36:0\|PC O-20:0_16:0 | 0.35737 | -1.4845 | 0.0026895 | 2.5703 |
| LPC 22:4 | 0.39121 | -1.354 | 0.041032 | 1.3869 |
| PC O-34:0\|PC O-18:0_16:0 | 0.40409 | -1.3073 | 0.0026236 | 2.5811 |
| LPS 16:1 | 0.42212 | -1.2443 | 0.012037 | 1.9195 |
| PC O-38:1\|PC O-16:0_22:1 | 0.44663 | -1.1629 | 0.0059525 | 2.2253 |
| PG 40:8\|PG 20:4_20:4 | 0.45459 | -1.1374 | 0.038696 | 1.4123 |
| LPS 22:3 | 0.45702 | -1.1297 | 0.0040539 | 2.3921 |
| PS O-38:2\|PS O-16:0_22:2 | 0.45959 | -1.1216 | 0.0029586 | 2.5289 |
| PS O-40:3\|PS O-18:0_22:3 | 0.4688 | -1.093 | 0.00037648 | 3.4243 |
| PC O-32:0\|PC O-16:0_16:0 | 0.4746 | -1.0752 | 0.00073911 | 3.1313 |
| PC 52:6\|PC 18:1_34:5 | 0.47727 | -1.0671 | 0.045052 | 1.3463 |
| PC 40:1\|PC 16:0_24:1 | 0.482 | -1.0529 | 0.026779 | 1.5722 |
| HexCer 40:1;2O\|HexCer 18:1;2O/22:0 | 0.48203 | -1.0528 | 0.0089678 | 2.0473 |
| LPE O-18:1 | 0.49607 | -1.0114 | 0.010121 | 1.9948 |
| NAGly 19:1;O | 2.6154 | 1.387 | 0.0012504 | 2.903 |
| NAGly 14:0 | 2.756 | 1.4626 | 0.0016359 | 2.7862 |
| PE 46:7\|PE 22:3_24:4 | 4.4711 | 2.1606 | 0.022109 | 1.6554 |

**Table 2.** DRP1-deficient NPCs have reduced ether-linked phospholipids and increased triglycerides (A) Fold change and p-values of significantly changed lipids between control and DRP1-deficient NPCs at day 8 of differentiation in positive ion mode. Decreased lipids in blue, increased lipids in red. **(B)** Fold change and p-values of significantly changed lipids between control and DRP1-deficient NPCs at day 8 of differentiation in negative ion mode. Decreased lipids in blue, increased lipids in red.

DRP1 Mutant vs Control – Positive Ion Mode
| Lipid Species | FC | log2(FC) | p-value | -log10(p) |
| --- | --- | --- | --- | --- |
| PC 52:2 | 0.0002754 | -11.826 | 2.65E-07 | 6.5765 |
| PC 28:1 | 0.018169 | -5.7824 | 1.93E-06 | 5.7141 |
| PC 27:1 | 0.040905 | -4.6116 | 0.01684 | 1.7736 |
| PC 25:0 | 0.064316 | -3.9587 | 0.026736 | 1.5729 |
| ST 27:2;O | 0.069839 | -3.8398 | 0.01619 | 1.7907 |
| PS 46:5 | 0.1132 | -3.1431 | 0.0059423 | 2.226 |
| PC 27:0 | 0.11601 | -3.1077 | 0.037041 | 1.4313 |
| PC 58:7 | 0.1555 | -2.685 | 0.0018642 | 2.7295 |
| PE 40:6\|PE 18:1_22:5 | 0.16178 | -2.6279 | 0.00014779 | 3.8304 |
| ST 27:0;O | 0.16772 | -2.5758 | 0.026172 | 1.5822 |
| PC 24:0 | 0.17549 | -2.5105 | 0.001076 | 2.9682 |
| PC 29:1 | 0.1894 | -2.4005 | 0.037444 | 1.4266 |
| LPC 16:1/0:0 | 0.18948 | -2.3999 | 4.83E-05 | 4.316 |
| PE 23:3\|PE 7:0_16:3 | 0.19339 | -2.3704 | 0.0061964 | 2.2079 |
| PC 32:4 | 0.21001 | -2.2515 | 0.019785 | 1.7037 |
| SM 34:0;2O | 2.6051 | 1.3813 | 0.016926 | 1.7715 |
| TG O-58:3\|TG O-18:0_18:1_22:2 | 2.6274 | 1.3936 | 0.048502 | 1.3142 |
| TG 64:6\|TG 18:1_24:2_22:3 | 2.6626 | 1.4128 | 0.036225 | 1.441 |
| TG O-52:2\|TG O-18:1_16:0_18:1 | 2.7218 | 1.4446 | 0.012732 | 1.8951 |
| TG O-54:2\|TG O-17:0_18:1_19:1 | 2.7302 | 1.449 | 0.023851 | 1.6225 |
| TG 64:5\|TG 18:1_20:2_26:2 | 2.8391 | 1.5055 | 0.026735 | 1.5729 |
| TG 56:2\|TG 18:0_18:1_20:1 | 2.9313 | 1.5515 | 0.014045 | 1.8525 |
| TG O-56:2\|TG O-18:0_18:1_20:1 | 2.9621 | 1.5666 | 0.02625 | 1.5809 |
| TG O-52:1\|TG O-18:0_16:0_18:1 | 2.9787 | 1.5747 | 0.0045663 | 2.3404 |
| TG 58:3\|TG 18:1_20:1_20:1_ID | 3.0549 | 1.6111 | 0.014379 | 1.8423 |
| TG 54:2;1O\|TG 18:1_18:1_18:0;1O | 3.0904 | 1.6278 | 0.043519 | 1.3613 |
| TG O-61:4\|TG O-21:1_18:1_22:2 | 3.1734 | 1.666 | 0.016544 | 1.7814 |
| TG 61:5;1O\|TG 20:1_20:1_21:3;1O | 3.2591 | 1.7045 | 0.02031 | 1.6923 |
| TG 62:3\|TG 18:1_20:1_24:1 | 3.3521 | 1.7451 | 0.042671 | 1.3699 |
| TG 60:3\|TG 16:0_18:1_26:2 | 3.8108 | 1.9301 | 0.015639 | 1.8058 |

DRP1 Mutant vs Control – Negative Ion Mode
| Lipid Species | FC | log2(FC) | raw.pval | -log10(p) |
| --- | --- | --- | --- | --- |
| PC 33:1;O\|PC 16:0_17:1;O | 0.0082594 | -6.9197 | 0.031756 | 1.4982 |
| PC 34:3;O\|PC 18:1_16:2;O | 0.04953 | -4.3355 | 0.035712 | 1.4472 |
| PC 26:1\|PC 12:0_14:1 | 0.05121 | -4.2874 | 0.0012348 | 2.9084 |
| PC 27:0\|PC 11:0_16:0 | 0.10151 | -3.3003 | 0.046507 | 1.3325 |
| PE 28:1\|PE 12:0_16:1 | 0.12046 | -3.0534 | 0.006128 | 2.2127 |
| PC 24:0\|PC 10:0_14:0 | 0.1384 | -2.8531 | 0.0021968 | 2.6582 |
| PE 28:0\|PE 14:0_14:0 | 0.1437 | -2.7989 | 0.0003883 | 3.4108 |
| NAGly 19:1;O | 0.1553 | -2.6869 | 0.029393 | 1.5318 |
| PE 33:1;2O\|PE 16:0_17:1;2O | 0.16627 | -2.5884 | 0.039451 | 1.4039 |
| PS 28:0\|PS 14:0_14:0 | 0.18212 | -2.4571 | 0.0076791 | 2.1147 |
| PC 28:1;O\|PC 14:1_14:0;O | 0.18432 | -2.4397 | 0.011042 | 1.957 |
| LPS 16:1 | 0.18705 | -2.4185 | 0.0002879 | 3.5406 |
| CL 60:2\|CL 14:0_16:1_14:0_16:1 | 0.18817 | -2.4099 | 0.0008735 | 3.0587 |
| PC 54:5\|PC 18:1_36:4 | 0.19302 | -2.3732 | 0.0099371 | 2.0027 |
| PC O-34:3\|PC O-16:2_18:1 | 0.20673 | -2.2742 | 0.020305 | 1.6924 |
| SM 34:0;2O | 2.3439 | 1.2289 | 0.026067 | 1.5839 |
| SHexCer 32:1;2O | 4.2452 | 2.0858 | 0.018332 | 1.7368 |

### PEX11β deficiency alters neural rosette morphology and neural progenitor pool abundance

Given the lipidomic alterations in PEX11β-deficient NPCs and the established importance of ether-linked phospholipids in neurodevelopment (Teigler et al., 2009), we revisited the effects of PEX11β deficiency on neural differentiation. Although neural identity was unchanged between control and PEX11β-deficient NPCs in two-dimensional culture, this system lacks the polarized architecture of the neuroepithelium. Therefore, we differentiated control and PEX11β KO iPSCs into neural rosettes, which recapitulate the apical-basal organization of neuroepithelial cells during early neural tube formation (Elkabetz et al., 2008; Wilson and Stice, 2006). The tight junction marker zonula occludens-1 (ZO-1) localizes apically and marks the central lumen, with PAX6+ neural progenitors extending radially from the lumen. The neural progenitor cells, which surround the lumen, have been shown to have similar identity to the cells found in the neural epithelial layer of the neural tube (Elkabetz et al., 2008). After 8 days of differentiation, neural rosettes were analyzed for identity and morphology, using PAX6 to quantify neural progenitor number and ZO-1 to measure lumen area **(Figure 8A)**. PEX11β KO neural rosettes exhibited a significantly increased number of PAX6+ neural progenitors and increased lumen area compared to control neural rosettes **(Figure 8B-8E, Supplementary Figures 7A-7C))**. To assess whether differences in apoptosis or proliferation accounted for this phenotype, neural rosettes were stained for cleaved poly (ADP-ribose) (clPARP), a hallmark of apoptosis, and Kiel 67 (Ki-67), a cell proliferation marker found exclusively in dividing cells (late G1, S, G2, and M phases). There were no differences in clPARP or Ki67 levels between control and PEX11β KO neural rosettes **(Figure 8C-8D)**. Thus, the enlarged lumen size and increased neural progenitor pool in PEX11β KO neural rosettes are unlikely to be attributable to differences in apoptosis or cell proliferation. Collectively, these findings show that PEX11β deficiency has functional consequences for early neural patterning.

**Figure 8.**
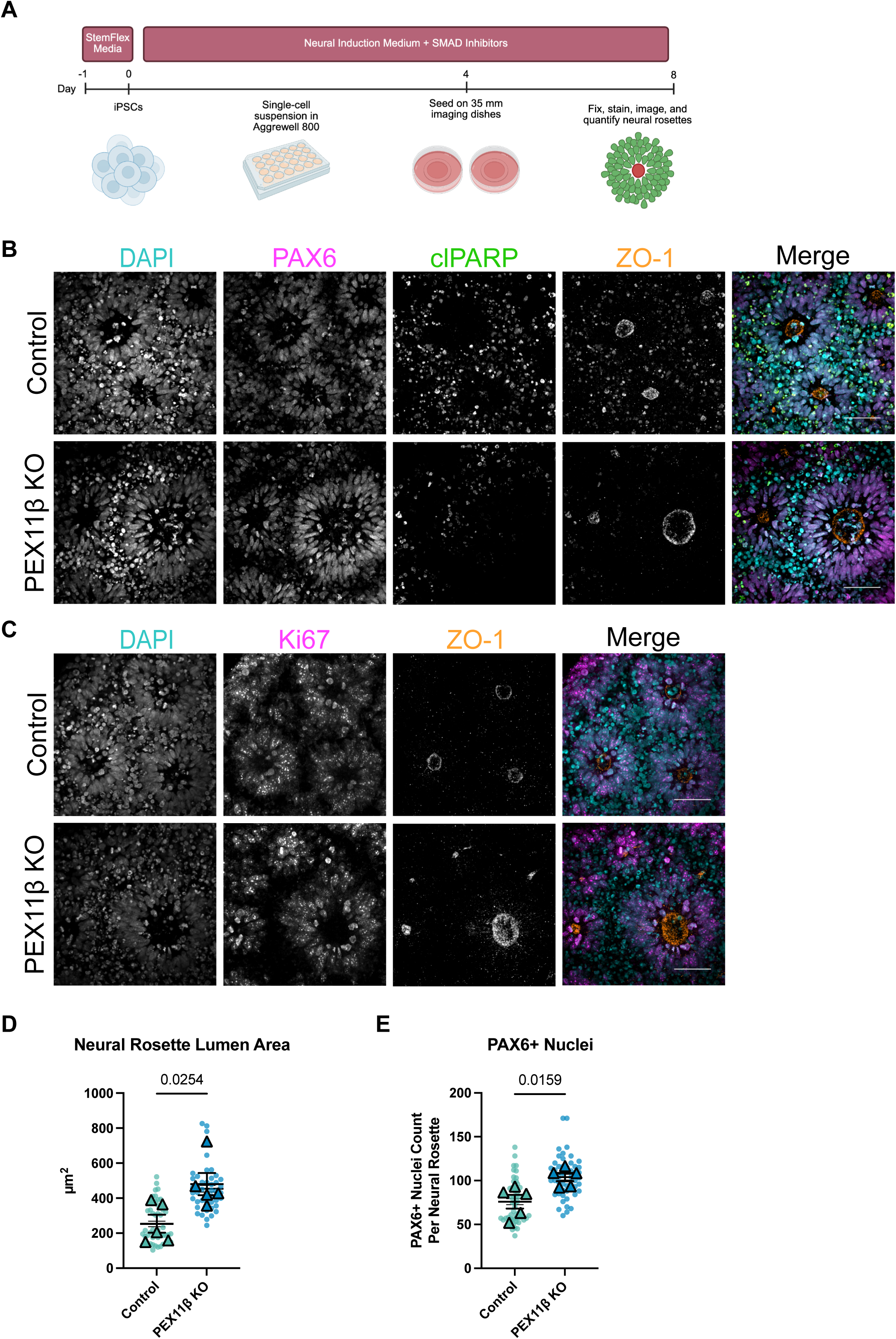
PEX11β KO NPCs neural rosettes have increased lumen area and neural progenitor number. **(A)** Schematic of neural rosette differentiation. **(B)** Representative maximum intensity projections of PAX6 (neural progenitors), clPARP (apoptosis), and ZO-1 (lumen) in control and PEX11β KO neural rosettes at day 8 of differentiation. Scale bars = 50μm (SDC). **(C)** Representative maximum intensity projections of Ki67 (cells in phases S, G2, or M) and ZO-1 (lumen) in control and PEX11β KO neural rosettes at day 8 of differentiation. Scale bars = 50μm (SDC). **(D)** Quantification of neural rosette lumen area (ZO-1) in control and PEX11β KO neural rosettes. Each dot represents the average per field of view (10 per n), each triangle represents a biological replicate (n=5). Analyzed using Welch’s t-test; error bars represent mean ± SEM. **(E)** Quantification of the number of PAX6+ neural progenitors per neural rosette in control and PEX11β KO neural rosettes. Each dot represents the average per field of view (10 per n), each triangle represents a biological replicate (n=5). Analyzed using Welch’s t-test; error bars represent mean ± SEM.

## Discussion

PEX11β is a central regulator of peroxisomal fission, from early membrane elongation to recruitment of MFF and FIS1, the receptors for DRP1, which mediates final membrane scission (Koch et al., 2004, 2003, 2010; Koch and Brocard, 2012). Individuals with PEX11β mutations present with neurological symptoms, including seizures and developmental delay (Ebberink et al., 2012; Henning et al., 2024; Khoddam et al., 2024; Malekzadeh et al., 2021; Taylor et al., 2017; Tian et al., 2019; Tsinopoulou et al., 2023), yet how PEX11β deficiency affects peroxisomal metabolism and cell fate during early human neurogenesis has remained unexplored. Given the established relationship between mitochondrial morphology and neural cell fate decisions during early neurogenesis (Iwata et al., 2020; Khacho et al., 2016), and the fact that peroxisomes are also highly dynamic organelles, we sought to examine whether peroxisomal morphology and metabolism similarly drive neural cell fate decisions during human neurogenesis. This study provides the first analysis of PEX11β deficiency in a human iPSC-derived neural system, with two goals: (i) to define the cell biology of PEX11β loss in human neural cells and (ii) to determine how peroxisomal elongation influences metabolism and early neural cell fate.

Consistent with prior reports, PEX11β KO iPSCs and NPCs displayed elongated peroxisomes. However, unlike most PEX11β-deficient systems, peroxisome number was unchanged in PEX11β-deficient iPSCs. During differentiation, peroxisomal morphology, abundance, and PEX11β expression were dynamically regulated in parallel with *PAX6* expression, suggesting a developmental transition from *de novo* biogenesis to fission-dependent remodeling. iPSCs contained numerous small peroxisomes, whereas NPCs exhibited fewer, larger organelles, consistent with shifting metabolic demands analogous to the glycolysis-to-OXPHOS transition in mitochondria (Agostini et al., 2016; Iwata et al., 2020; Khacho et al., 2016; Zheng et al., 2016) These findings indicate that peroxisomal morphology is developmentally tuned during neural commitment.

PEX11β-deficient NPCs showed reduced peroxisome fission frequency and reduced colocalization between peroxisomes and the downstream components of the peroxisomal fission machinery. These results highlight the critical role of PEX11β in peroxisomal fission. Because PEX11β has been reported to also participate in peroxisomal membrane elongation, the basis of the elongated peroxisomal morphology in PEX11β-deficient cells remains unresolved. A prevailing model attributes this phenotype to lipid flow from the ER to peroxisomes, which is mediated by the VAPB-ACBD5 tether (Costello et al., 2017; Hua et al., 2017). However, disrupting this tether did not shorten peroxisomes in PEX11β-deficient NPCs; instead, ACBD5 knockdown increased peroxisomal surface area and volume in both PEX11β-deficient and control NPCs. These results point to a mechanism where peroxisomal elongation in NPCs is largely independent of ER lipid flow. We propose that continued import of peroxisome matrix and membrane proteins in the absence of efficient fission deforms the peroxisomal membrane, leading to elongation, and that altered lipid composition following ER-peroxisome tether disruption may further affect recruitment of fission factors.

Following characterization of peroxisomal morphology and dynamics, we confirmed that PEX11β deficiency did not result in changes in mitochondrial morphology in iPSCs, nor changes in mitochondrial morphology or respiration in iPSC-derived NPCs. Hallmark studies examining the link between mitochondrial morphology and neural cell fate have often relied on genetic or pharmacological manipulation of DRP1, which is critical for both mitochondrial and peroxisomal fission. However, the effects of perturbing DRP1 on peroxisomal fission and metabolic function have remained largely unexplored in such studies. Therefore, our PEX11β-deficient iPSC and iPSC-derived neural model systems, which do not have changes in mitochondrial morphology or respiration, offer a means to study the independent contributions of elongated peroxisomal morphology on metabolism and cell fate decisions during early human neurogenesis.

Studies in mouse models have linked Pex11β loss to neural migration defects, impaired neural differentiation, and neural cell death (Ahlemeyer et al., 2012; Esmaeili et al., 2016; Li et al., 2002), yet these phenotypes are more severe than those observed in individuals with PEX11β mutations, who typically show normal brain structure by MRI (Ebberink et al., 2012; Taylor et al., 2017). However, in other neurodevelopmental disorders, it has been shown that alterations in early transcriptional programs during early neurogenesis can, in turn, alter cell fate decisions in neural progenitor cells to change neurodevelopmental trajectories, resulting in consequences later in development (Jourdon et al., 2023). Additionally, human neurodevelopment differs markedly from mouse neurodevelopment, with expanded radial glial pools, greater interneuron diversity, and more complex circuitry (Molnár et al., 2019). Therefore, exploring PEX11β-deficiency during early neurogenesis in human systems is critical, as the neurological symptoms observed in individuals deficient in PEX11β may be a result of early cell fate dysregulation at the neural progenitor stage.

PEX11β-deficient NPCs showed reduced *TUBB3* expression at day 8 of differentiation, reminiscent of other fission-impaired systems that bias progenitors toward self-renewal (Iwata et al., 2020; Khacho et al., 2016). However, by day 12 of NPC differentiation, there were no longer differences in neural identity in PEX11β-deficient NPCs, suggesting recovery from this imbalance or a compensatory mechanism. Lipidomic analysis of PEX11β-deficient NPCs at day 8 of differentiation revealed a reduction in ether-linked phospholipids and an increase in a sphingomyelin species with a very-long-chain fatty acid side chain. This suggests perturbations in peroxisomal metabolic functions in PEX11β-deficient NPCs, potentially pointing to deficiencies in the plasmalogen synthesis and β-oxidation pathways, for which peroxisomes are essential. In polarized neural rosettes, PEX11β deficiency produced enlarged lumens and expanded PAX6+ pools, consistent with altered symmetric–asymmetric divisions of the neural progenitor pool (Elkabetz et al., 2008). Progenitor imbalances have also been found in Pex11β-deficient epithelia due to spindle misorientation (Asare et al., 2017), though whether this mechanism operates in PEX11β-deficient human iPSC-derived neural rosettes remains unclear.

Overall, our results link PEX11β deficiency in neural cells to changes in peroxisomal metabolism, and early alterations in neural fate in human neurogenesis. Phenotypes were milder than in mouse models, suggesting species-specific sensitivity, yet early progenitor imbalance could still reshape developmental later trajectories without overt malformations. Several key open questions remain: 1) What is the main driver of peroxisomal elongation in PEX11β-deficient systems? 2) What specific peroxisomal functions contribute to the neurodevelopmental phenotypes in PEX11β deficiency? 3) What are the effects of the early neurodevelopmental perturbations observed in our systems on the rest of the neurodevelopment trajectory?

Ultimately, leveraging this PEX11β-deficient iPSC system for differentiation into more developmentally mature human brain models, coupled with metabolic profiling in these systems, provides a powerful platform for elucidating the specific cell-fate changes that may contribute to neurological symptoms in individuals with PEX11β mutations. Eventually, these systems may serve as tools for testing potential therapeutic options.

### Limitations

A limitation of this study is that PEX11β deficiency results in reduction in peroxisome number, which creates difficulty when interpreting whether findings are the result of elongated peroxisomal morphology or reduced peroxisome number. We propose that the overall content of peroxisomes (i.e. peroxisome biogenesis proteins, matrix enzymes) in PEX11β-deficient peroxisomes may be similar to those in control peroxisomes, although this requires further validation. We also found that in PEX11β-deficient iPSCs, peroxisome number is not changed, offering a platform for investigating the morphological effects of PEX11β deficiency in a system where peroxisome number is not affected.

We were also keen to quantify peroxisomal fission events, as these dynamics have not been explored live in neural systems. An important caveat of our peroxisomal event localizer approach is that it relies on segmentation and spatial resolution, and some events scored as fission may not be true fission events, but rather peroxisomes in proximity that later move apart. To mitigate this limitation, peroxisomes were analyzed in 3D and fission events were manually verified, confirming a reduction in fission events in PEX11β KO NPCs compared to control NPCs.

Another limitation of this study is that the CRISPR-generated PEX11β knockout represents complete loss of function, whereas some affected individuals harbor hypomorphic or missense variants that may retain partial activity. Full knockout could therefore exaggerate some phenotypes or fail to capture allele-specific mechanisms relevant to human disease.

Finally, the differentiation systems used in this study model early stages of human neurogenesis. In the future, complementing these studies with more developmentally mature model systems, such as iPSC-derived neural organoids, will be important to define the impact of PEX11β deficiency on later developmental processes.

### Methods Cell lines

The PGP-1 (GM2338) iPSC line was used in this study. The GM2338 (RRID: CVCL_F182) line is from a 55-year-old healthy, Caucasian male. The iPSCs were maintained on Matrigel-coated plates (Corning #354277) and cultured in StemFlex medium (Thermo Fisher Scientific #A3349401) at 37°C with 5% CO2. Cells were checked for differentiation and cell culture medium was changed daily. Every 3-4 days, the cells were passaged. For passaging, iPSCs were dissociated with Gentle Cell Dissociation Reagent (Stem Cell Technologies #100-0485) and incubated at room temperature for 4 minutes. All experiments were performed under the supervision of the Vanderbilt Institutional Human Pluripotent Cell Research Oversight (VIHPCRO) Committee. Cells were checked for contamination periodically, validated for pluripotency using the PluriTest Global Gene Expression Profile Assay (Thermo Fisher Scientific #A38154) and karyotyped every 20 passages.

### Single-cell flow sorting

PEX11β knockout (KO) iPSC pools were obtained from Synthego. Two hours prior to sorting, the media on the PEX11β KO iPSC pool was changed to StemFlex medium (Thermo Fisher Scientific #A3349401) supplemented with 10μM of Y-27632 Rho/Rock pathway inhibitor (Stem Cell Technologies #72307). One hour prior to sorting, 96-well plates (Corning #3596) were coated with Matrigel (Corning #354277). Thirty minutes before sorting, Cloning Medium (StemFlex + 10% CloneR [StemCell Technologies #05888] + 1% Penicillin Streptomycin [Thermo Fisher Scientific #15-140-122]) was warmed to 37°C. To harvest cells for sorting, cells were first washed with DBPS (Thermo Fisher Scientific #14190144), followed by dissociation with Gentle Cell Dissociation Reagent (Stem Cell Technologies #100-0485) for 8 minutes at 37°C. After removal of Gentle Cell Dissociation Reagent, StemFlex + 10μM of Y-27632 Rho/Rock pathway inhibitor was added, and the cells were gently scraped for harvesting. The cells were then transferred to a 15mL conical and centrifuged at 200x*g* for 2 minutes. After the supernatant was aspirated, the cells were resuspended in StemFlex + CloneR. 100μL/well of Cloning Medium was then added to the Matrigel-coated 96-well plates. Resuspended cells were passed through a 40μm filter (Pluriselect #435004051) immediately before sorting. Cells were single cell sorted using a 100μm nozzle, with 1 cell per well. Once sorting was complete, the plates were spun down at 120x*g* for 3 minutes to attach the cells to the bottom of the plate. The cells were incubated for 72 hours, at which point a full media change was performed with Cloning Medium. 96 hours after sorting, a full media change was performed with StemFlex medium without CloneR. Wells with a detectable iPSC colony were marked, while any wells containing more than 1 colony were avoided. Media changes were continued until the colonies were ready to passage (∼12 days after sorting). Colonies were passaged using the iPSC passaging procedures described above.

### Sample preparation and LC-MS for analysis of PEX11β

Coomassie-stained gel regions were excised and diced into 1mm^3^ cubes. Gel pieces were equilibrated in 100mM ammonium bicarbonate, reduced with 4.5 mM DTT at 56°C for 20 minutes, and alkylated with 10mM iodoacetamide for 30 minutes, protected from light (Tateishi et al., 2024). Gels were destained with 1:1 (v/v) 100% acetonitrile and 50mM ammonium bicarbonate and were dehydrated by addition of 100% acetonitrile. Following removal of acetonitrile, gel pieces were dried and in-gel digested with trypsin. Trypsin Gold (mass spectrometry grade, Promega #V5280) was diluted in 25mM ammonium bicarbonate (pH 8.0) to 10ng/μL and added to cover the dehydrated gel pieces, on ice. After 20 minutes, 25mM ammonium bicarbonate (10μL) was added to the gel pieces, and digestion was performed by incubation at 37°C overnight. The resulting tryptic peptides were recovered by 2 sequential extractions (15 minutes each) with 100μL of 60% acetonitrile/0.1% trifluoroacetic acid. Extracts were dried, reconstituted in 0.2% aqueous formic acid, and analyzed by LC-coupled tandem mass spectrometry (LC-MS/MS). Using a Dionex Ultimate 3000 nanoLC and autosampler, peptides were loaded onto a reverse phase analytical column (360µm O.D. x 100µm I.D.), packed with 20cm of C18 material as described previously (Tateishi et al., 2024). Peptides were gradient-eluted at a flow rate of 350nL/minute, using a 90-minute gradient. The mobile phase solvents consisted of 0.1% formic acid/ 99.9% water (solvent A) and 0.1% formic acid/99.9% acetonitrile (solvent B). The gradient was as follows: 2–38% B in 75 minutes; 38–90% B in 7 minutes; 90% B for 1 minutes; 90–2% B in 1 minutes; 2% B for 6 minutes. Peptides were analyzed using a data-dependent method on a Thermo Scientific Orbitrap Exploris 240 mass spectrometer, equipped with a nanoelectrospray source. The instrument method consisted of MS1 using an AGC target value of 3 x 106, followed by up to 20 MS/MS scans using an AGC target of 1x105. The intensity threshold for MS/MS was set to 1 x 104, HCD collision energy was 30 normal collision energy (nce), and dynamic exclusion (15 seconds) was enabled. For identification of peptides, tandem mass spectra were searched with Sequest (Thermo Fisher Scientific) against a *Homo sapiens* database created from the UniprotKB protein database (www.uniprot.org). Variable modification of +15.9949 on Met (oxidation) and +57.0214 on Cys (carbamidomethylation) were included for database searching. Search results were assembled using Scaffold 5.1.0. (Proteome Software), using a minimum filtering criteria of 95% peptide and protein probability. Peptides were also analyzed via data-independent acquisition (DIA) on a Bruker timsTOF HT mass spectrometer. Using a nanoElute2 LC system (Bruker), samples were loaded onto a reverse phase column, packed with 20cm of C18 material. The column was connected with a 20µm CaptiveSpray emitter, and peptides were gradient-eluted at 500nL/minute using a 40-minute gradient (2-35% B in 30 minutes). diaPASEF data were acquired using 12 windows (ramps), spanning 0.75–1.3 V-s/cm2 1/k0 with an accumulation time of 100 ms. Spectronaut, version 20, (Biognosys) (Bruderer et al., 2015) was used for analysis of DIA data. A directDIA search was performed with default settings for identification, which included a false discovery rate (FDR) of 1% for peptides and protein groups. The precursor Qvalue and the experiment-wide protein Qvalue cutoffs were set to 0.01. Method evaluation was added in the experimental settings for the DIA analysis workflow.

### Trilineage differentiation

Trilineage differentiation was performed according to the manufacturer’s protocol (Stem Cell Technologies #05230). Briefly, when iPSCs reached 90% confluency, they were dissociated with Gentle Cell Dissociation Reagent (Stem Cell Technologies #100-0485) and incubated at 37°C with 5% CO2 for 8 minutes. They were then resuspended in StemFlex medium with 10μM Y-27632 Rho/Rock pathway inhibitor (Stem Cell Technologies #72307), scraped gently with cell lifters, and centrifuged at 300*xg* for 5 minutes. Cells were plated onto 12-well plates at 800,000 cells/well for the endoderm and ectoderm differentiations, and 200,000 cells/well for the mesoderm differentiation. 24 hours later, media was changed to the lineage-specific media. Cells for mesoderm and endoderm differentiation were maintained at 37°C and 5% CO2 for 5 days, while cells for ectoderm differentiation were maintained at 37°C and 5% CO2 for 7 days. Media was changed daily.

### Differentiation of iPSCs into neural progenitors

Neural progenitor cells (NPCs) were generated from iPSCs using dual SMAD neural induction using 10μM of SB431542 (Reprocell #04-0010-10) and 1μM dorsomorphin (Sigma Aldrich #P5499-5MG). When iPSCs reached 80–90% confluency, they were dissociated with Gentle Cell Dissociation Reagent (Stem Cell Technologies #100-0485) and incubated at 37°C with 5% CO2 for 8 minutes. DMEM/F12 (Thermo Fisher Scientific #11320033) with 10μM Y-27632 Rho/Rock pathway inhibitor (Stem Cell Technologies #72307) was then added to the cells, which were scraped with cell lifters and centrifuged at 300*xg* for 5 minutes. They were then resuspended in essential 6 medium (Lippmann et al., 2015, 2014) containing 10μM Y-27632 Rho/Rock pathway inhibitor (Stem Cell Technologies #72307) and SMAD inhibitors SB4 and dorsomorphin, single-celled further with a P1000 pipette, and plated at 2.0–2.5x10^6^ cells/well of a 6-well plate or 1.0x10^6^ cells/well of a 12-well plate. NPCs were incubated at 37°C and media was changed daily. At day 8 of differentiation, NPCs were collected for downstream processing.

### Differentiation of iPSCs into neural rosettes

When iPSCs reached 80–90% confluency, they were dissociated using Gentle Cell Dissociation Reagent and incubated at 37°C with 5% CO2 for 8 minutes. DMEM/F12 (Thermo Fisher Scientific #11320033) with 10μM Y-27632 Rho/Rock pathway inhibitor (Stem Cell Technologies #72307) was next added to the cells, and the cells were detached with cell lifters prior to centrifugation at 300*x*g for 5 minutes. Cells were then resuspended in STEMdiff Neural Induction Medium (Stem Cell Technologies #05839) with 10μM Y-27632 Rho/Rock pathway inhibitor, further dissociated with a P1000 pipette, and plated at 3x10^6^ cells/well in an AggreWell 800 plate (Stem Cell Technologies #34815) pre-treated with 500μL of Anti-Adherence Rinsing Solution for 5 minutes (Stem Cell Technologies #07010). Embryoid bodies were incubated at 37°C with 5% CO2, with minimal disruption, for 48 hours. A 50% media change was performed with STEMdiff Neural Induction Medium on day 2. On day 4, embryoid bodies were harvested using a 40μm mesh cell strainer (Pluriselect #435004051) and transferred to 2, 35mm imaging dishes (Cellvis #D35-10-1.5-N) coated with Matrigel. Media changes were performed daily and the cells were collected at day 8 of differentiation.

### RNAi transfection

Commercially available siRNA targeting *ACBD5* (Thermo Fisher Scientific #s40666) was used to transiently knock down *ACBD5* in control and PEX11β KO NPCs. NPCs were generated as described above, and medium was changed on day 1 of neural differentiation. Cells were transfected per manufacturer protocol using Lipofectamine RNAiMax (Thermo Fisher Scientific #13778150) in Essential 6 medium on days 2 and 3 of neural differentiation. Silencer Select Negative Control No. 1 (Thermo Fisher Scientific #4390844) was used as a control. Cells were collected for downstream analysis at day 4 of differentiation.

### RNA extraction and cDNA synthesis

Cells were scraped into 1000μL of Trizol reagent (Thermo Fisher Scientific #15596018) and flash frozen. After thawing, the samples were allowed to sit for 5 minutes at room temperature. Then, 200μL of chloroform was added (Sigma-Aldrich #C2432-4X25ML) and the samples incubated at room temperature for 2–3 minutes. The samples were then centrifuged at 12,000x*g* for 15 minutes at 4°C, followed by collection of the aqueous phase. Next, 500μL of isopropanol (Sigma-Aldrich #I9516-500ML) was added to precipitate RNA. The samples were incubated for 25 minutes at room temperature, followed by centrifugation at 12,000x*g* for 10 minutes at 4°C. The RNA pellet was washed with 75% ethanol, semi-dried, and resuspended in 30μL of nuclease-free water (Thermo Fisher Scientific #AM9937). Following quantification and adjustment of the volume of all samples to 1μg/μL, the samples were treated with DNAse (New England Biolabs #M0303), which was inactivated using EDTA (Sigma-Aldrich #324506-100ML). cDNA was generated using 10μL of this volume using the manufacturer’s protocol (Thermo Fisher Scientific #4368814).

### Quantitative RT-PCR (RT-qPCR)

RT-qPCR was run on the QuantStudio 3 Real-Time PCR machine according to manufacturer instructions with 1μg of cDNA sample, SYBR Green Universal Master Mix (Thermo Fisher Scientific #4364346), and the primers outlined in the Key Resources Table.

### Western blot

Cells were collected in lysis buffer containing PhosSTOP (Roche #04906837001), Protease Inhibitor Cocktail (PIC) (Roche #04693116001), and PMSF (RPI #P20270-10.0) diluted in 1% Triton-X100 (Sigma-Aldrich #T9284) in PBS. Cells were vortexed every 5 minutes for 30 minutes, followed by sonication for further cell lysis. 50μg of protein were loaded and run on 4–20% Mini-Protean TGX precast protein gels (Bio-Rad #4561094) in Tris-Glycine-SDS buffer. Gels were transferred onto polyvinylidene difluoride membranes (PVDF) (Sigma Aldrich #IEVH00005) at 0.31 amps for 1 hour at room temperature. After transfer, the membrane was blocked in 5% milk (RPI #M17200500) in TBST for 1 hour at room temperature on a benchtop rocker. The primary antibodies were diluted in 5% milk and incubated overnight at 4°C on a benchtop rocker. Blots were washed 3 times for 5 minutes with TBST, followed by incubation with HRP-conjugated secondary antibodies in 5% milk for 1 hour at room temperature on a benchtop rocker. Blots were developed with ECL Plus Reagent (Thermo Fisher Scientific #32106) or SuperSignal West Femto (#Thermo Fisher Scientific #34095). They were then imaged on a CCD AI600 chemiluminescent imager. Band density was quantified using Fiji (imagej.net/software/fiji/). Data were normalized to the loading control, and data were transformed and normalized to the average of the biological replicates of the scramble control.

### Immunofluorescence

Cells were fixed with 4% paraformaldehyde in 1X PBS for 15 minutes at room temperature, followed by permeabilization with 1% Triton X-100 (Sigma-Aldrich #T9284) in 1X PBS for 5 minutes at room temperature. Cells were then washed 3 times with PBS, blocked in 10% BSA in 1X PBS, and incubated overnight at 4°C with primary antibody diluted in 10% BSA (Key Resources Table). Cells were washed 3 times with PBS followed by incubation with secondary antibody (Key Resources Table) diluted in 10% BSA for 1 hour at room temperature. Following secondary antibody incubation, cells were washed an additional 3 times with 1X PBS, followed by incubation with Hoechst (1mg/mL) (Thermo Fisher Scientific #H3570) in 1X PBS for 10 minutes at room temperature. Cells were washed once with 1X PBS and sealed with Fluoromount-G slide mounting medium (Thermo Fisher Scientific #00-4958-02). For MitoTracker experiments, cells were incubated with MitoTracker Red CMXRos (Thermo Fisher Scientific #M46752) at a concentration of 100nm for 30 minutes at 37°C and 5% CO2 prior to fixation.

Neural rosettes were fixed on day 8 of differentiation with 100% ice-cold methanol (Thermo Fisher Scientific #A454-4) for 10 minutes at -20°C. Next, they were washed with TBS and blocked at room temperature with 5% donkey serum (Jackson ImmunoResearch #017-000-121) + 0.3% Triton X-100 in tris-buffered saline (TBS), followed by overnight incubation at 4°C with primary antibodies diluted in blocking solution (Key Resources Table). After being washed three times with 1X TBS, cells were incubated with secondary antibody in 5% donkey serum + 0.3% Triton X-100 in TBS for 2 hours. After secondary antibody incubation, cells were washed an additional 3 times with 1X TBS and then incubated with Hoechst (1mg/mL) in TBS for 10 minutes at room temperature. After 1 additional wash, samples were mounted in Fluoromount-G slide mounting medium.

### Live imaging of peroxisomal morphology

Neural progenitors were generated from iPSCs as described above and plated onto Matrigel-coated 35mm imaging dishes (Cellvis #D35-10-1.5-N). At day 3 of neural progenitor differentiation, CellLight Peroxisome-GFP, BacMam 2.0 (Thermo Fisher Scientific #C10604) was added to cells at 5 particles per cell, and the cells were incubated at 37°C and 5% CO2. After 16 hours, Peroxisome Cell Light was removed and the cells washed once with essential 6 medium. Prior to live cell imaging, cells were switched into FluoroBrite DMEM (Thermo Fisher Scientific #A1896701) with NucBlue (Thermo Fisher Scientific #R37605).

### Image acquisition

For imaging of stem cell identity, trilineage differentiation, neural progenitor identity, and neural rosettes, samples were imaged on the Nikon W1 Spinning Disc Confocal with a 20X Plan Apo 0.75NA objective, 60X Plan Apo 1.40 NA oil objective, or 100X Plan Apo 1.45 NA oil objective and a prime 95B CMOS camera. For live imaging of peroxisomal morphology, cells were imaged on the Nikon W1 Spinning Disc Confocal with a 100X Plan Apo 1.49 NA oil objective and a prime 95B CMOS camera. Cells were imaged in a temperature-controlled chamber maintained at 37°C and 5% CO2. Super-resolution images of peroxisomal morphology, mitochondrial morphology, and colocalization of peroxisomal antibodies with antibodies marking peroxisomal fission proteins were acquired using a Nikon Super Resolution by Optical Pixel Reassignment (SoRa) microscope equipped with a 100X Plan Apo 1.45 NA oil objective and a Hamamatsu ORCA-Fusion BT camera. Imaging of peroxisomal morphology across neural differentiation was performed using a Nikon Structured Illumination Microscope (SIM) equipped with a 100X Apo TIRF 1.49 NA WD 0.12 oil objective and an Andor iXon Ultra DU-897 EMCCD monochrome camera. All imaging was done with at least 3 biological replicates and 7 images per replicate.

### Image analysis

Quantification of mitochondrial and peroxisomal morphology was performed in NIS-Elements version 5.42.06 (Nikon). Peroxisomes or mitochondria were segmented in 3D and data were extracted from the resulting 3D mask. Major axis length, surface area, volume, and count measurements were exported into Excel. Counts of DAPI, counts of transcription factors, and measurement of neural rosette lumen area were performed manually using Fiji (imagej.net/software/fiji/).

### Peroxisomal event localizer (PEL) analysis

A total of 31 cells were analyzed: n1 = 5 control and 4 PEX11β KO cells; n2 = 6 control and 5 KO cells; and n3 = 5 control and 6 KO cells. Time-lapse 3D z-stack imaging was conducted over ∼180 timepoints with 10 z-slices every 5 seconds at a 0.1µm step size and 100ms exposure. To assess potential photobleaching effects on measurements, analyses were performed on three temporal windows: (1) full acquisition (all ∼180 timepoints), (2) first 50 timepoints, and (3) first 100 timepoints. Automated event detection was performed using custom ImageJ macros implementing the Mitochondrial Event Localizer (MEL) (Theart et al., 2020), repurposed here as the peroxisomal event localizer (PEL). Importantly, MEL/PEL is organelle-agnostic: it detects topological split and merge events in 3D instance-labeled organelle masks (i.e., uniquely labeled 3D connected components) and does not rely on mitochondria-specific assumptions. The code is available for download at https://github.com/rensutheart/MEL-Fiji-Plugin. For each timepoint, peroxisomal structures were segmented from the 3D z-stack and converted to uniquely labeled 3D connected components (instance labeling) and tracked across sequential frames. Fission events were identified when a single labelled structure at time *t* split into two (or more) daughter structures at *t+1* and remained split at *t+2* (a conservative persistence criterion to suppress transient segmentation artifacts). Movies were not drift-registered, as peroxisomes are near-diffraction-limited and robust registration was unreliable; observed drift was gradual and MEL/PEL detects events via changes in instance topology rather than absolute position. Peroxisome counts at each timepoint were recorded for normalization. Fission frequency was calculated as the number of fission events per peroxisome per timepoint. To account for the hierarchical structure of the data and avoid pseudoreplication, statistical comparisons followed a two-stage approach. First, per-cell metrics were aggregated to compute replicate-level means, calculated separately for control and KO genotypes within each biological replicate. Second, paired statistical tests compared KO versus control using these replicate means (n = 3 biological replicates), treating each biological replicate as the unit of analysis. Paired t-tests assessed normally distributed differences.

### Seahorse Mito Stress Test

Neural progenitor cells were plated onto Seahorse XF97 V3 PS cell culture microplates 2 days before the assay (day 6 of neural differentiation) at 80,000 cells per well. One hour before the assay, cell medium was changed to XF DMEM (Agilent #103575-100) containing 1mM pyruvate (Agilent #103578-100), 2mM L-glutamine (Agilent #103579-100), and 10mM D-glucose (Agilent #103577100). Oxygen consumption rate (OCR) was measured sequentially after addition of 1μM oligomycin, 0.5μM FCCP, and 0.5μM rotenone plus antimycin A (all included in Agilent #103015-100).

### Liquid chromatography-mass spectrometry for untargeted lipidomics

To prepare samples, a one-phase method was used to extract lipids. Briefly, 0.5mL of methanol/methyl tert-butyl ether/chloroform mix (1.3:1:1) was added to a frozen pellet of NPCs (2x10^6^ cells total), spiked with 10μL EquiSPLASH-lipidomics internal standard mix (Avanti Research), briefly vortexed and shaken gently for 20 minutes, and centrifuged at 20,000*xg* for 15 minutes at 10°C. The supernatant was transferred to a clean Eppendorf tube, evaporated under a gentle stream of N2 gas, resuspended in 100μL methanol/chloroform (9:1), and 2μL were used for LC-HRMS (high-resolution MS) analysis. Each sample was injected two times—one injection in positive electrospray ionization mode, followed by one in negative mode. Pooled QCs were injected to assess the performance of the LC and MS instruments at the beginning, in the middle and at the end of each sequence. Discovery lipidomics data were acquired using a Vanquish UHPLC (ultrahigh performance liquid chromatography) system interfaced to a Q Exactive HF quadrupole/orbitrap mass spectrometer (Thermo Fisher Scientific).

Chromatographic separation was performed with a reverse-phase Acquity BEH C18 column (1.7μm, 2.1x150mm, Waters, Milford, MA) at a flow rate of 250μl/min and 50°C. Mobile phases were made up of 10 mM ammonium formate and 0.1% formic acid in (A) water/acetonitrile (40:60) and in (B) acetonitrile/isopropyl alcohol (10:90). Gradient conditions were as follows: 0–1 minutes, B = 20%; 1–8 minutes, B = 20–100%; 8–10 minutes, B = 100%; 10–10.5 minutes, B = 100–20 %; 10.5–15 minutes, B = 20%. The total chromatographic run time was 15 minutes. Mass spectra were acquired over a precursor ion scan range of m/z 200 to 1,600 at a resolving power of 60,000 using the following HESI-II source parameters: spray voltage 4kV (3kV in negative mode); capillary temperature 250°C; S-lens RF level 60V; N2 sheath gas 40; N2 auxiliary gas 10; auxiliary gas temperature 350°C. MS/MS spectra were acquired for the most abundant precursor ions (Top7) with an AGC target of 1x10^5^, a maximum injection time of 100 ms, and a normalized collision energy of 15, 30, 40. High resolution mass spectrometry data were processed with MS-DIAL version 4.90 in lipidomics mode. MS1 and MS2 tolerances were set to 0.01 and 0.025 Da respectively. Minimum peak height was set to 30,000 to decrease the number of false positive hits. Peaks were aligned with RT tolerance of 0.1 minutes, and mass tolerance of 0.015 Da. Default lipid library was used, solvent type was set to ammonium formate (HCOONH4) to match the solvent used for separation, and the identification score cut off was set to 80%. All lipid classes were made available for the search. After lipid annotation was completed, MS-DIAL results were exported into Excel and cleaned using minimum RSD for QC samples set to 20% and minimum ratio of QC to Blank set to 10. For species identified in both KO and controls, mean levels (areas under peak curves) and their variance were analyzed in Excel. Normalized data were uploaded to MetaboAnalyst for analysis (https://www.metaboanalyst.ca/).

### Statistical analysis

Experiments were performed with a minimum of 3 biological replicates and 2 technical replicates (2 control clones and 2 PEX11β KO clones). Normality of data was determined using the Shapiro-Wilk test. Statistical significance was determined using a Welch’s t-test, unless otherwise noted in the figure legend. GraphPad Prism v10 was used for all statical analysis and data visualization. Error bars in all bar graphs represent standard error of the mean unless otherwise indicated in the figure. Outliers were removed using the robust regression and outlier removal (ROUT) method where appropriate. For all statistical analyses, a statistically significant difference was established when P < 0.05.

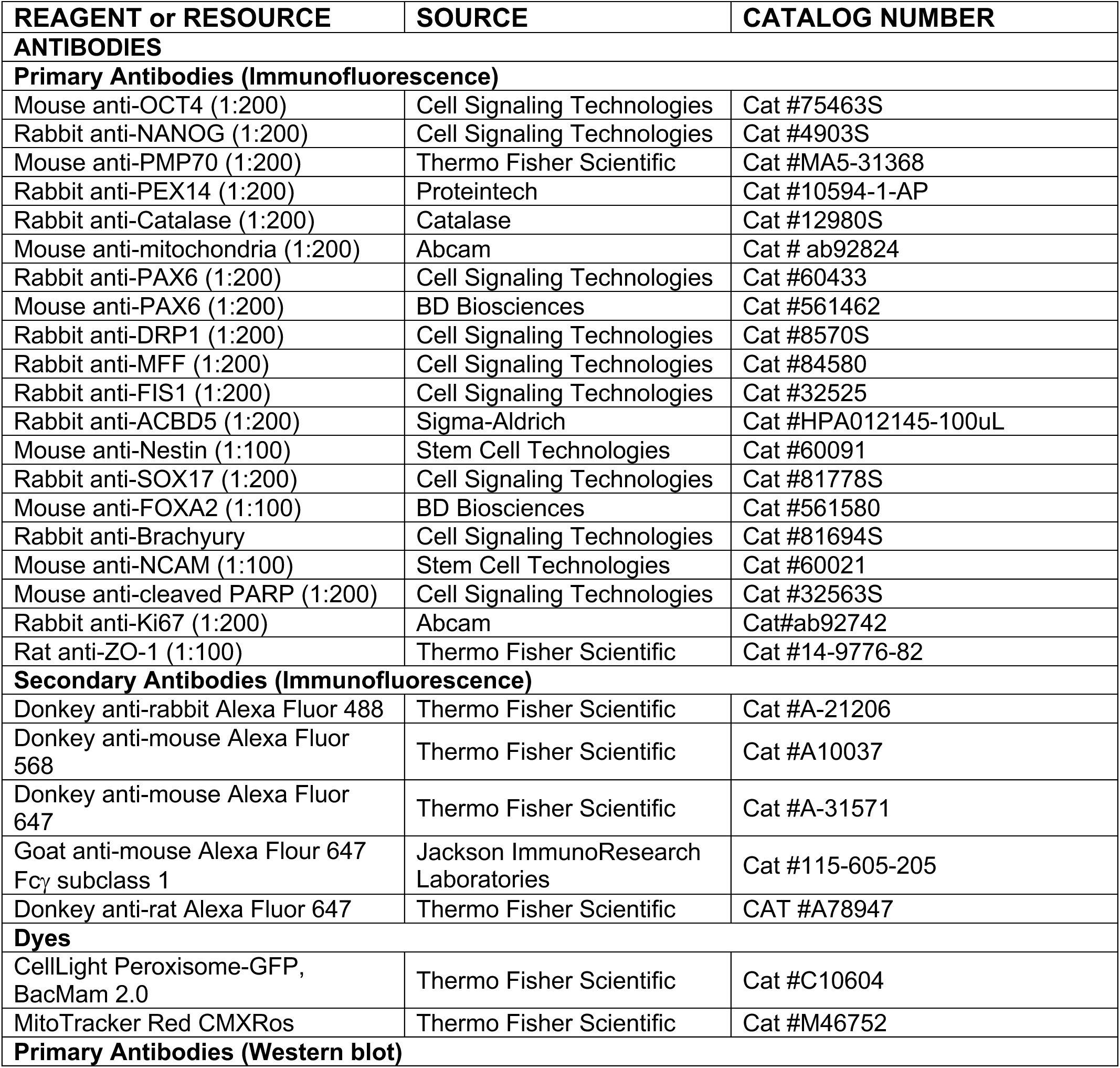
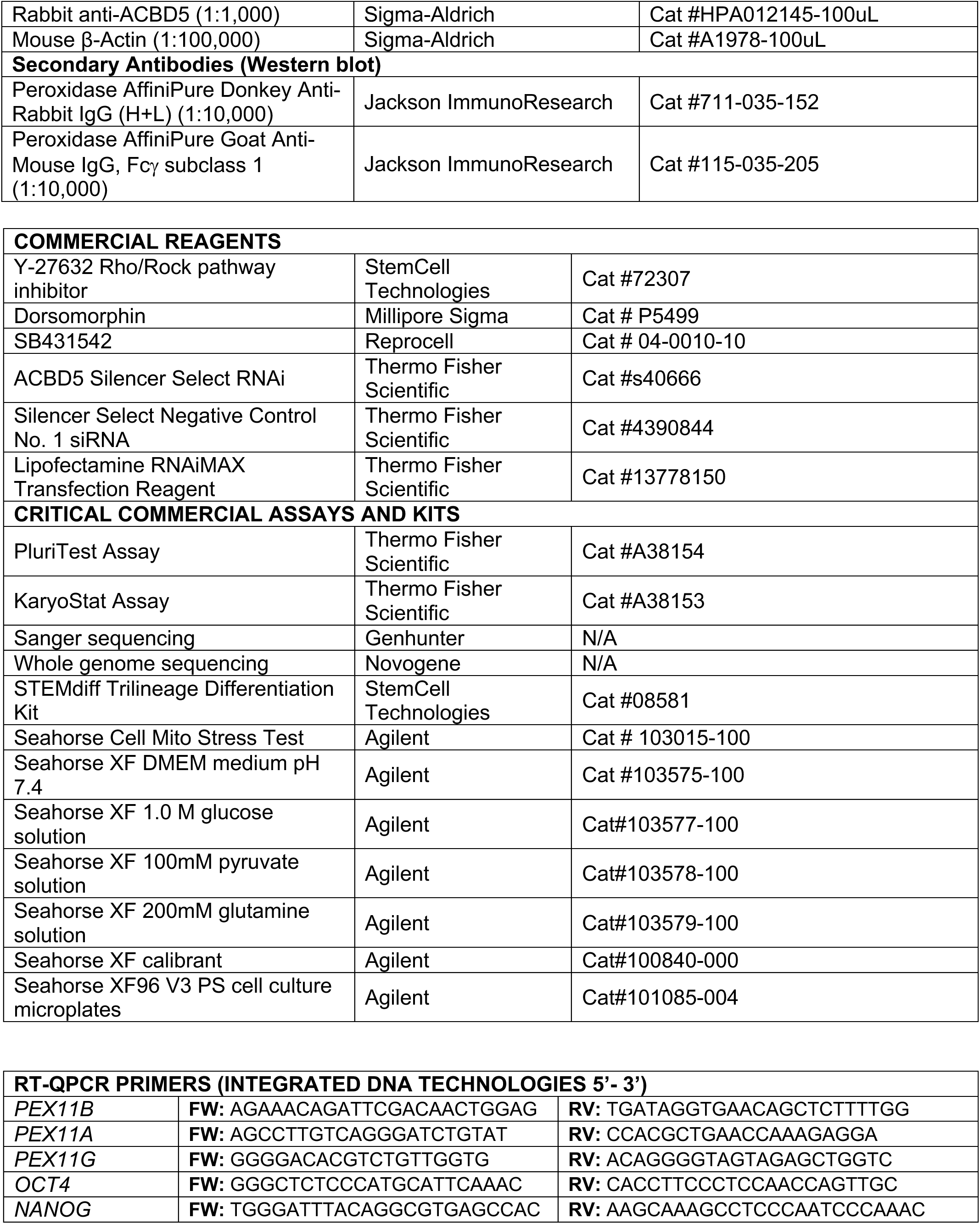
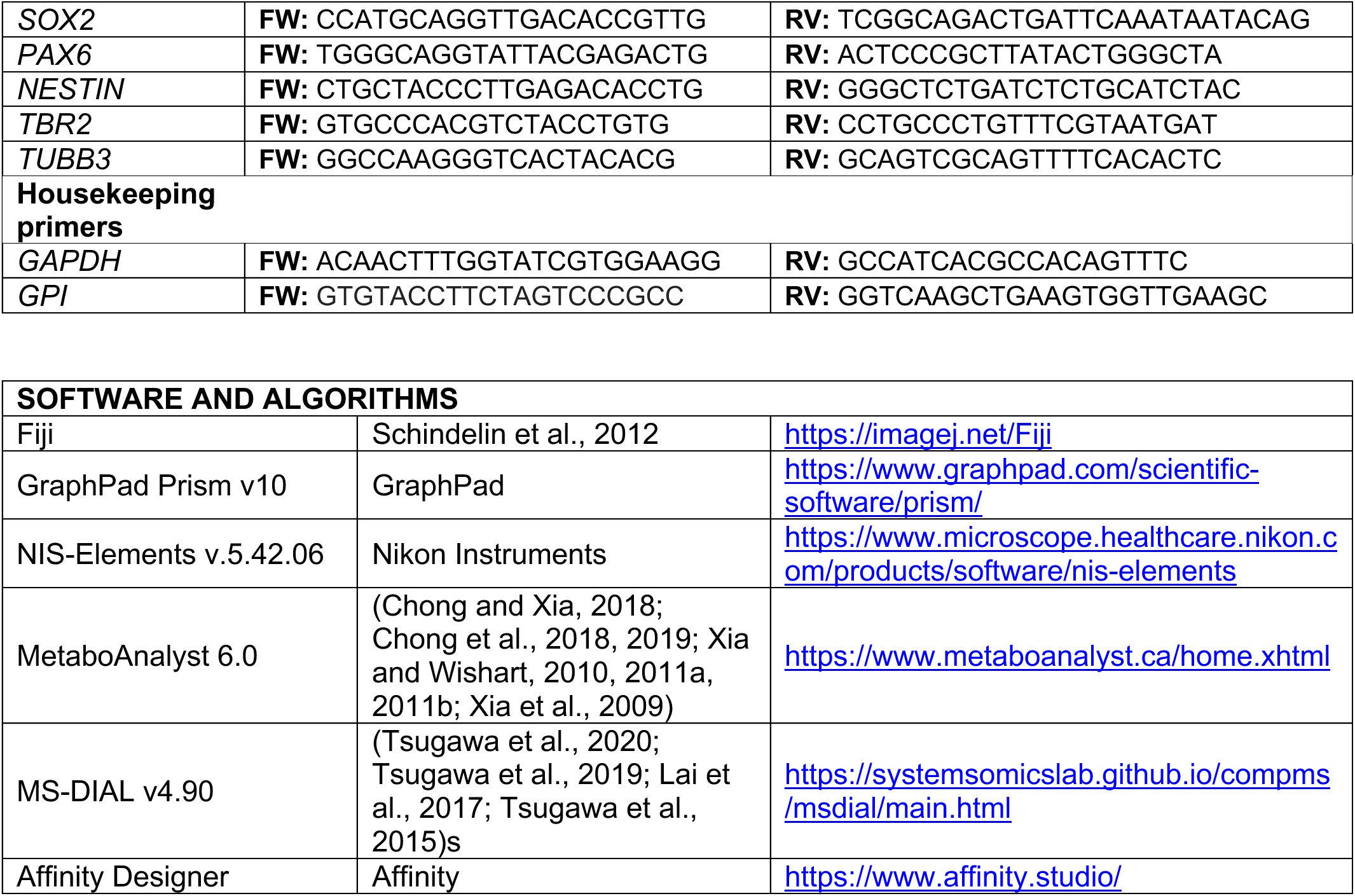

## Acknowledgements

We thank Gabriella Robertson and Caleb Hayes for their technical expertise and assistance with experimental design and interpretation. We thank Drs. Matthew Tyska, Kristopher Burkewitz, Kevin Ess, and Elma Zaganjor for their critical feedback during this project. We thank Dr. Kristie Rose in the Vanderbilt Mass Spectrometry Research Center for mass spectrometry analysis of PEX11β peptides and Dr. Sergei Chetyrkin in the Vanderbilt Mass Spectrometry Core Lab for untargeted lipidomics data collection. We would also like to thank Kari Seedle for providing advice and expertise with high-resolution microscopy and image analysis.

This work was supported by 2R35GM128915-06 (VG), a Vanderbilt Seeding Success Grant (VG), and 1F31HD114431-01A1 (CB). All Nikon SIM and SoRa microscopy was performed using the Vanderbilt Cell Imaging Shared Resource, supported by NIH grants CA68485, DK20593, DK58404, DK59637, EY08126, and 1S10MH130456-01A1. Flow Cytometry experiments were performed in the VMC Flow Cytometry Shared Resource. The VMC Flow Cytometry Shared Resource is supported by the Vanderbilt Ingram Cancer Center (P30 CA68485) and the Vanderbilt Digestive Disease Research Center (DK058404). Figures 1A, 2A, 7A, and 8A in the manuscript were created using Biorender.com.

