## Supplemental Figures and Legends for "Modeling the cell biology of PEX11β deficiency during human neurogenesis"

**Supplemental Figure Legends**

**Supplementary Figure 1. Validation of PEX11β KO iPSCs. (A)** qRT-PCR analysis of the PEX11 isoforms, PEX11β, PEX11α, and PEX11γ, in control and PEX11β KO iPSCs relative to control and normalized to two housekeeping genes (*GPI* and *GAPDH)*. Each dot on the graph represents a biological replicate; different colors represent different clones (n=3). Analyzed using Welch's t-test; error bars represent mean ± SEM. **(B)** Whole genome sequencing results confirming exonic, frameshift deletions in two PEX11β KO clones, resulting in early stop codons (denoted by fs*30 or fs*31). **(C)** Mass spectrometry analysis showing 40% sequence coverage of the PEX11β sequence in control iPSCs, with no detection of peptides matching PEX11β in PEX11β KO iPSCs. (**D)** Karyotyping indicates that control and PEX11β KO iPSC clones have a normal number and structure of chromosomes.

**Supplementary Figure 2. PEX11β KO iPSCs maintain stem cell identity. (A)** PluriTest score showing that both control and PEX11β KO iPSCs share a similar pluripotency signature compared to a non-iPSC control. Pluripotency score (y-axis) indicates how strongly the model-based pluripotency signature is expressed in the analyzed samples. Novelty score (x-axis) shows how well the transcriptional profile of the cells of interest matches the expected transcriptional profile. **(B)** qRT-PCR analysis of stem cell identity markers *OCT4*, *NANOG*, and *SOX2* in control and PEX11β KO iPSCs relative to control and normalized to two housekeeping genes (*GPI* and *GAPDH)*. Each dot on the graph represents a biological replicate; different colors represent different clones (n=3). Analyzed using Welch's t-test; error bars represent mean ± SEM. **(C)** Representative maximum intensity projections of peroxisomal morphology (PEX14) and mitochondrial morphology in control and PEX11β KO iPSCs. Scale bars = 10μm (SoRa).

**Supplementary Figure 3. PEX11β KO iPSCs differentiate into the three germ layers. (A)** Representative maximum intensity projections of control and PEX11β KO iPSCs differentiated into endoderm (SOX17, FOXA2). Scale bars = 100μm (SDC). **(B)** Representative maximum intensity projections of control and PEX11β KO iPSCs differentiated into mesoderm (BRACHYURY, NCAM). Scale bars = 100μm (SDC). **(C)** Representative maximum intensity projections of control and PEX11β KO iPSCs differentiated into ectoderm (PAX6, NESTIN). Scale bars = 100μm (SDC).

**Supplementary Figure 4. Peroxisome number is reduced in PEX11β-deficient NPCs but not iPSCs. (A)** Representative maximum intensity projections of peroxisomal morphology (PEX14) in control and PEX11β KO iPSCs. Scale bars = 10μm. Zoom scale bars = 2.5μm (SoRa). **(B)** Quantification of peroxisomal count in control and PEX11β KO iPSCs normalized to DAPI. Each dot represents the average per field of view (7 per n), each triangle represents a biological replicate, and different colors represent different clones (n=3). Analyzed using Welch's t-test; error bars represent mean ± SEM. **(C)** Representative maximum intensity projections of peroxisomal morphology (PEX14) in control and PEX11β KO NPCs at day 8 of differentiation. Scale bars = 10μm. Zoom scale bars = 2.5μm (SoRa). **(D)** Quantification of peroxisomal count in control and PEX11β KO NPCs at day 8 of differentiation, normalized to DAPI. Each dot represents the average per field of view (7 per n), each triangle represents a biological replicate, and different colors represent different clones (n=3). Analyzed using Welch's t-test; error bars represent mean ± SEM.

**Supplementary Figure 5. Peroxisomal fission is reduced in PEX11β KO NPCs. (A)** Representative still images from live 3D time-lapse imaging of peroxisomes in control NPCs at day 4 of differentiation using Peroxisome Cell Light (n=3, 5 cells per replicate). Cells were imaged at 5-second intervals for 15 minutes (final minute excluded from analysis due to photobleaching). Magenta arrows show points of observed peroxisomal fission. PEL overlay is rendered on frame 1 to indicate which structure undergoes fission in frame 2 and is maintained for an additional frame. Scale bars = 10μm (SDC). **(B)** Still images from live imaging of peroxisomes in PEX11β KO NPCs using Peroxisome Cell Light (n=3, 5 cells per replicate). Cells were imaged at 5-second intervals for 15 minutes (final minute excluded from analysis due to photobleaching). Magenta arrows show points of observed peroxisomal fission. PEL overlay is rendered on frame 1 to indicate which structure undergoes fission in frame 2, with fission maintained for the subsequent frame. Scale bars = 10µm (SDC). **(C)** Quantification of peroxisome fission events in the first 100 frames of peroxisome live imaging in control and PEX11β KO NPCs. Each dot on the graph represents a biological replicate. Analyzed using paired t-test; error bars represent mean ± SEM.

**Supplementary Figure 6. Colocalization of MFF, FIS1, and DRP1 with PMP70 in NPCs. A)** Representative maximum intensity projections of MFF, mitochondria (MitoTracker), and peroxisomes (PMP70) in control and PEX11β KO NPCs at day 8 of differentiation. Scale bars = 10µm (SoRa). **(B)** Representative maximum intensity projections of FIS1, mitochondria (MitoTracker), and peroxisomes (PMP70) (SoRa) in control and PEX11β KO NPCs at day 8 of differentiation. Scale bars = 10µm. **(C)** Representative maximum intensity projections of DRP1, mitochondria (MitoTracker), and peroxisomes (PMP70) in control and PEX11β KO NPCs at day 8 of differentiation. Scale bars = 10µm (SoRa).

**Supplementary Figure 7. Control and PEX11β KO neural rosettes. A-C)** Representative maximum intensity projections of PAX6 (neural progenitors) and ZO-1 (lumen) in control and PEX11β KO neural rosettes at day 8 of differentiation. Scale bars = 100µm (SDC).

**Video S1, related to Figure 5. Peroxisomal fission events in control NPCs.** Peroxisomal event localizer detection of peroxisomal fission events (red) in control NPCs. 3D time-lapse acquisition every 5 seconds for 15 minutes (final minute excluded due to photobleaching). Scale bars = 10µm (SDC).

**Video S2, related to Figure 5. Peroxisomal fission events in PEX11β KO NPCs.** Peroxisomal event localizer detection of peroxisomal fission events (red) in PEX11β KO

NPCs. 3D time-lapse acquisition every 5 seconds for 15 minutes (final minute excluded due to photobleaching). Scale bars = 10μm (SDC).

Supplementary Figure 1

**A**

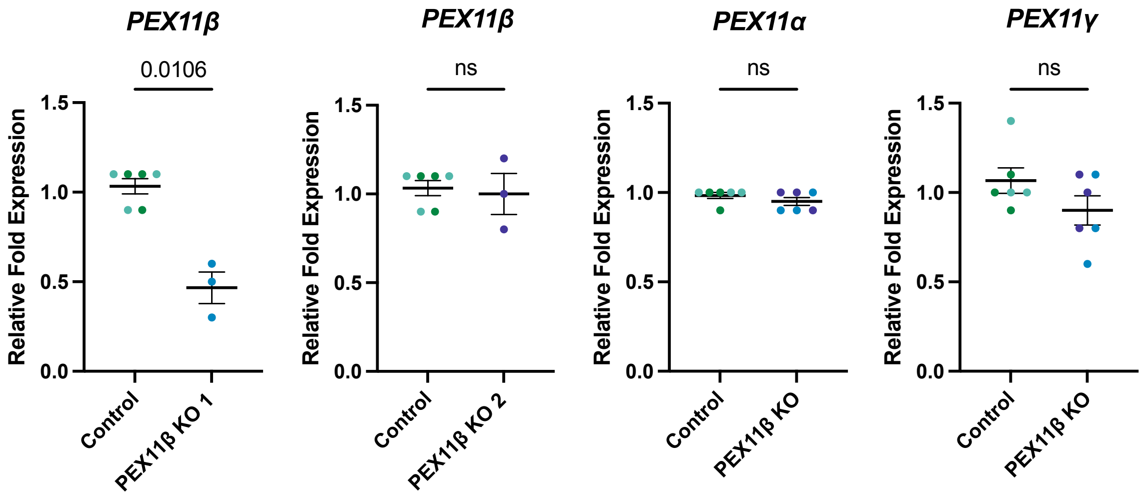


**B**

Key
PEX11β KO 1
PEX11β KO 2

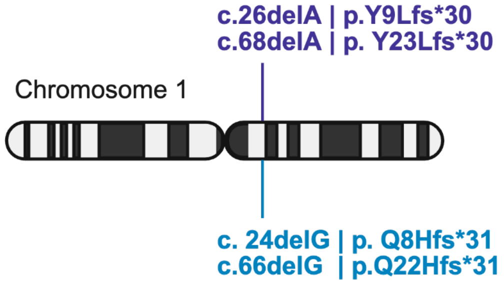


| | Priority | Function | ExonicFunc | Exon Position |
|---|---|---|---|---|
| PEX11β KO 1 | High | Exonic | Frameshift deletion | Exon 2 |
| PEX11β KO 2 | High | Exonic | Frameshift deletion | Exon 2 |

**C**

Control
PEX11β KO 1
PEX11β KO 2

**PEX11B – Human**

MDAWVRFSAQSQARERLCRAAQYACSLLGHALQRHGASPELQKQIRQLESHLSLGR
KLLRLGNSADALESAKRAVHLSDVVLRFCITVSHLNRALYFACDNVLWAGKSGLAPR
VDQEKWAQRSFRYYLFSLIMNLSRDAYEIRLLMEQESSACSRRLKGSGGGVPGGSE
TGGLGGPGTPGGGLPQLALKLRLQVLLLARVLRGHPPLLLDVVRNACDLFIPLDKLG
LWRCGPGIVGLCGLVSSILSILTLIYPWLRLKP

**D**

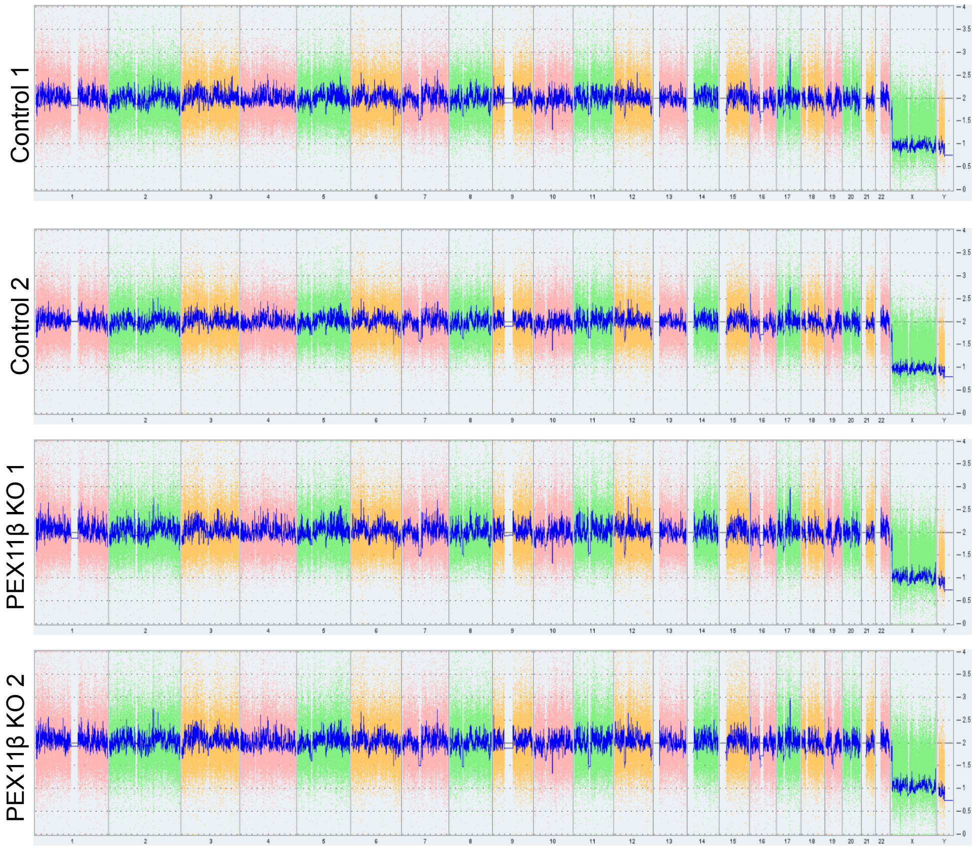

Supplementary Figure 2

**A**

| Sample | PluriTest Result | Pluripotency Score | Novelty Score |
|---|---|---|---|
| Control 1 | Pass | 40.2785 | 1.595706 |
| Control 2 | Pass | 41.47454 | 1.577171 |
| PEX11β KO 1 | Pass | 38.52407 | 1.620435 |
| PEX11β KO 2 | Pass | 39.58702 | 1.570617 |
| iPSC Control | Pass | 37.90687 | 1.229926 |
| Non-iPSC Control | Fail | -45.36237 | 2.718222 |

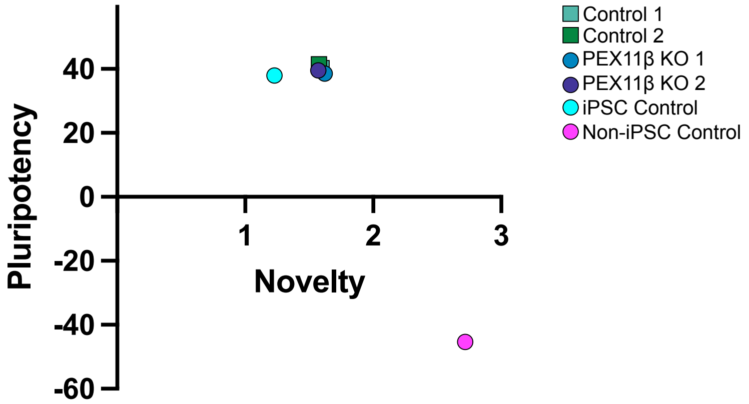


**B**

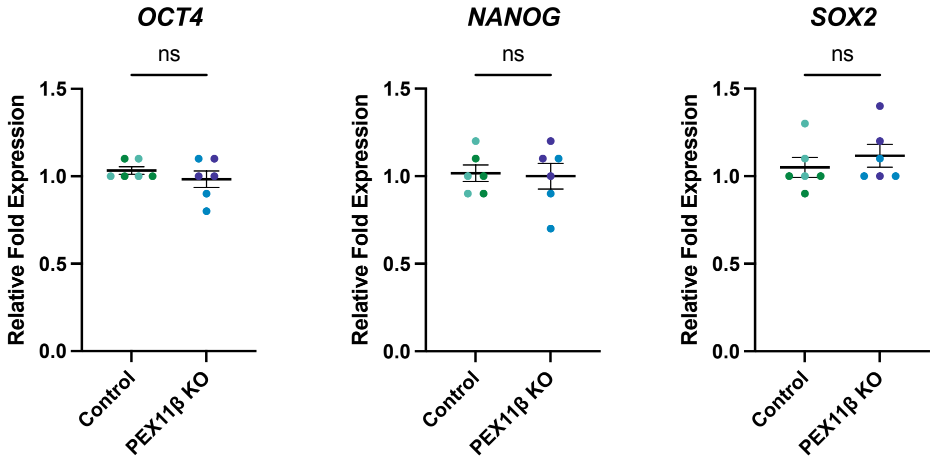


**C**

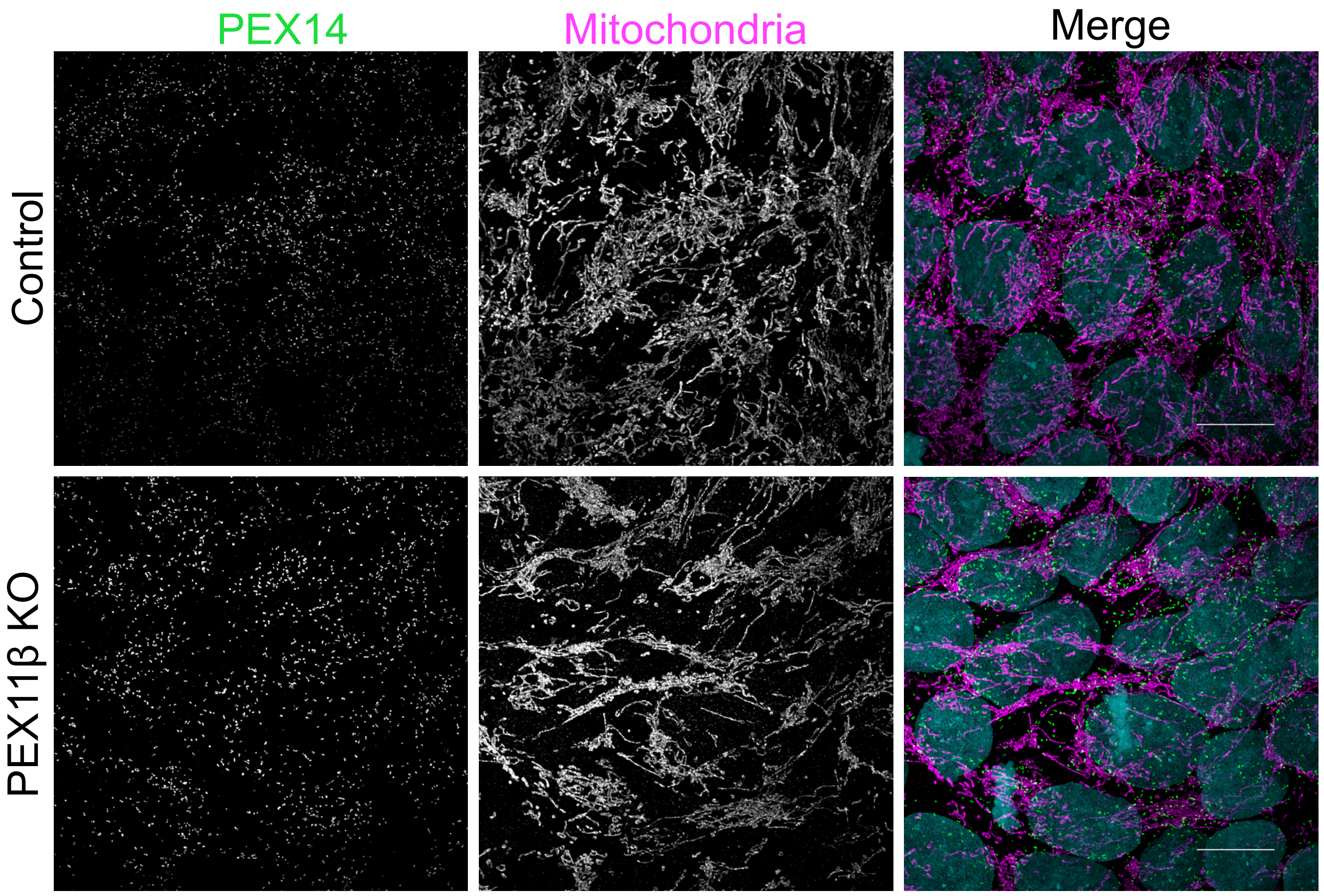

Supplementary Figure 3

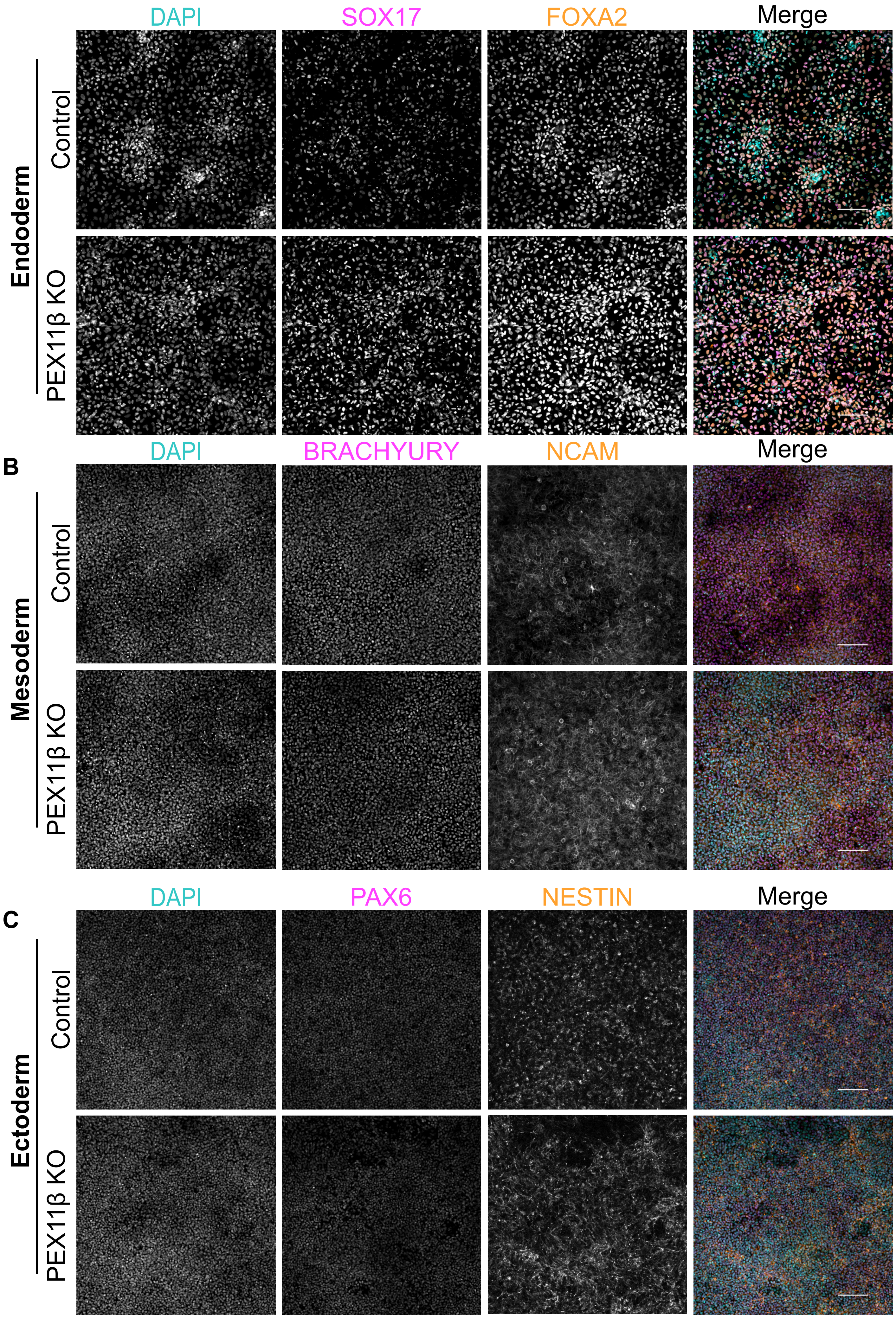

Supplementary Figure 4

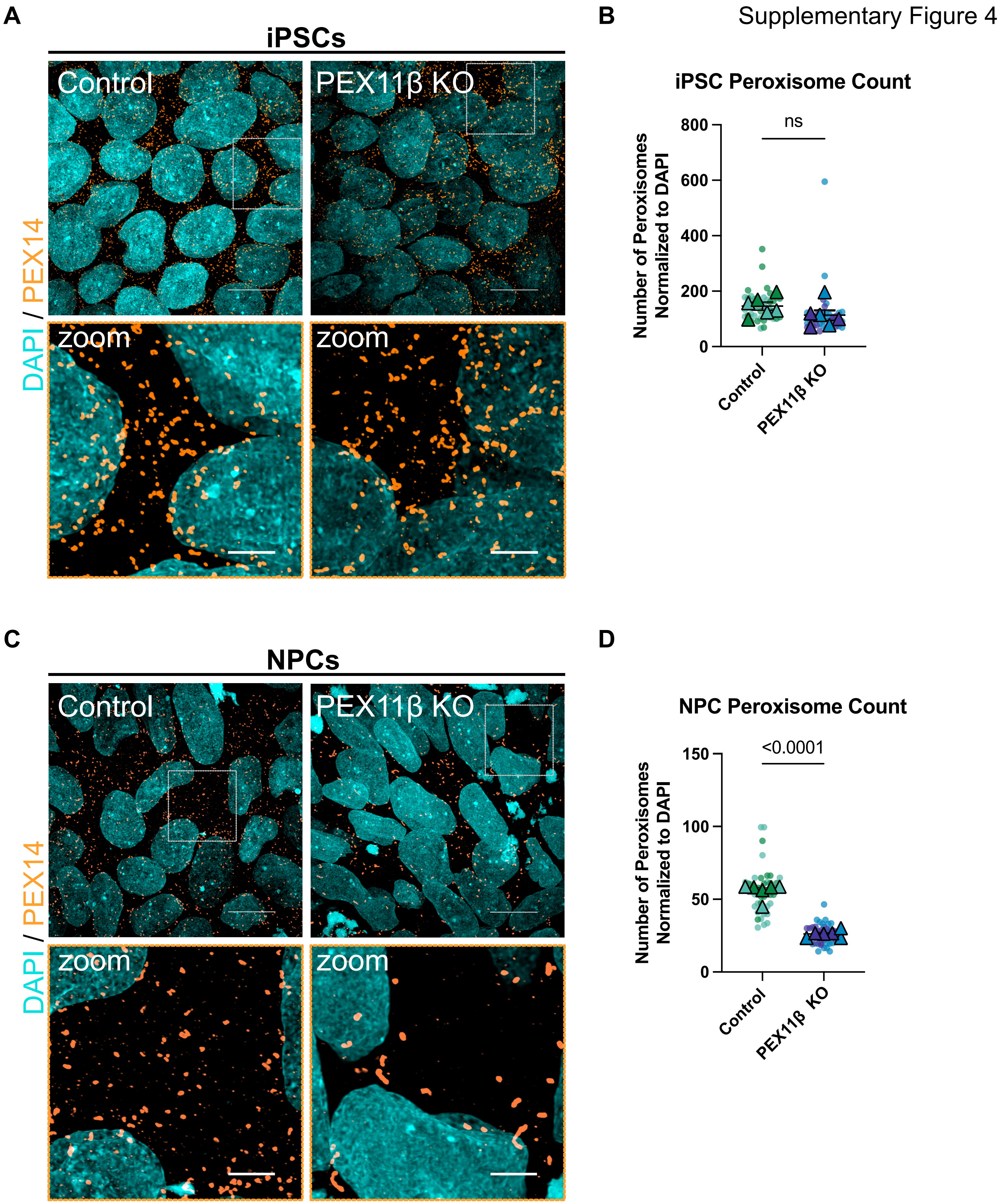

Supplementary Figure 5

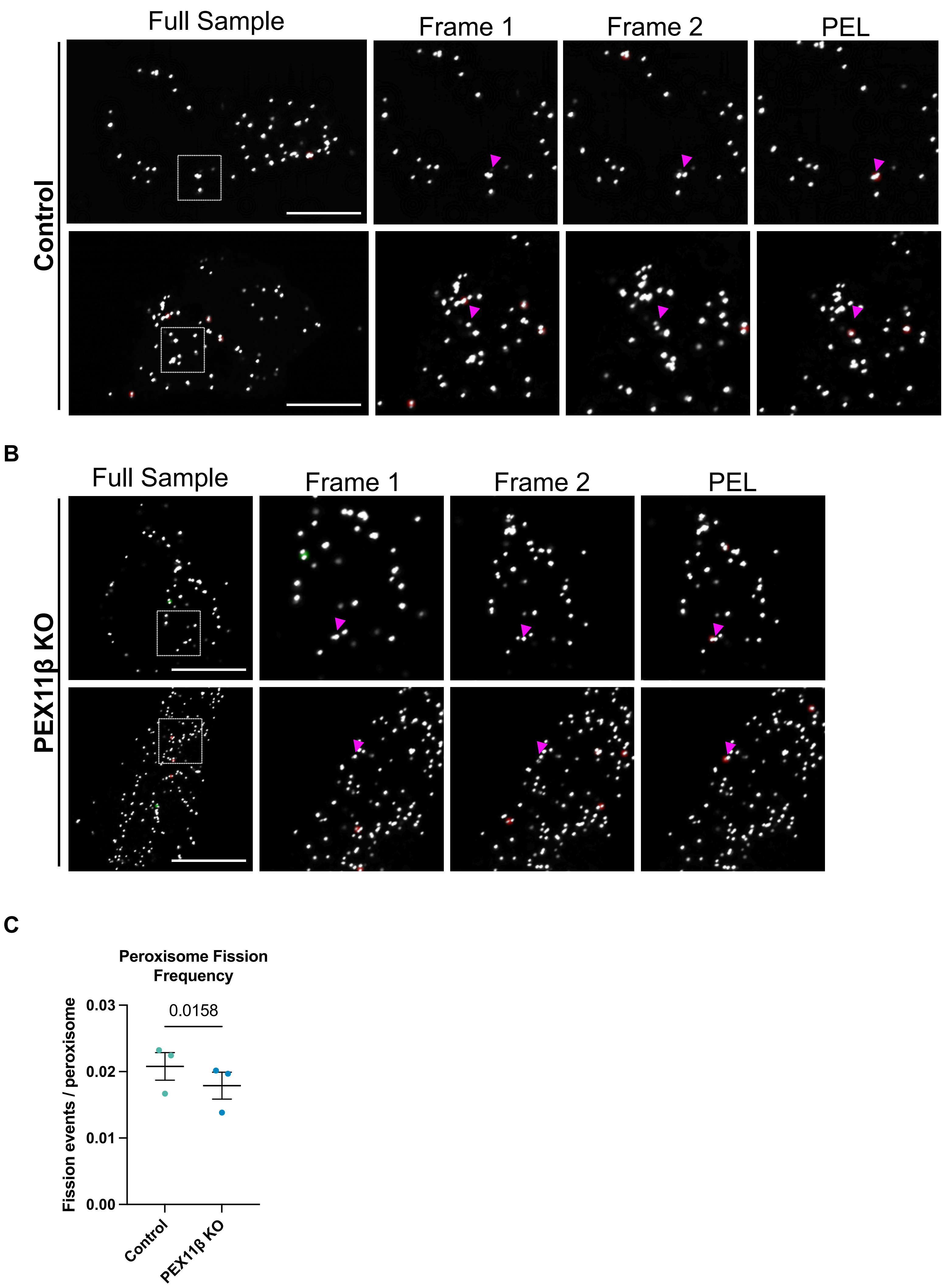

Supplementary Figure 6

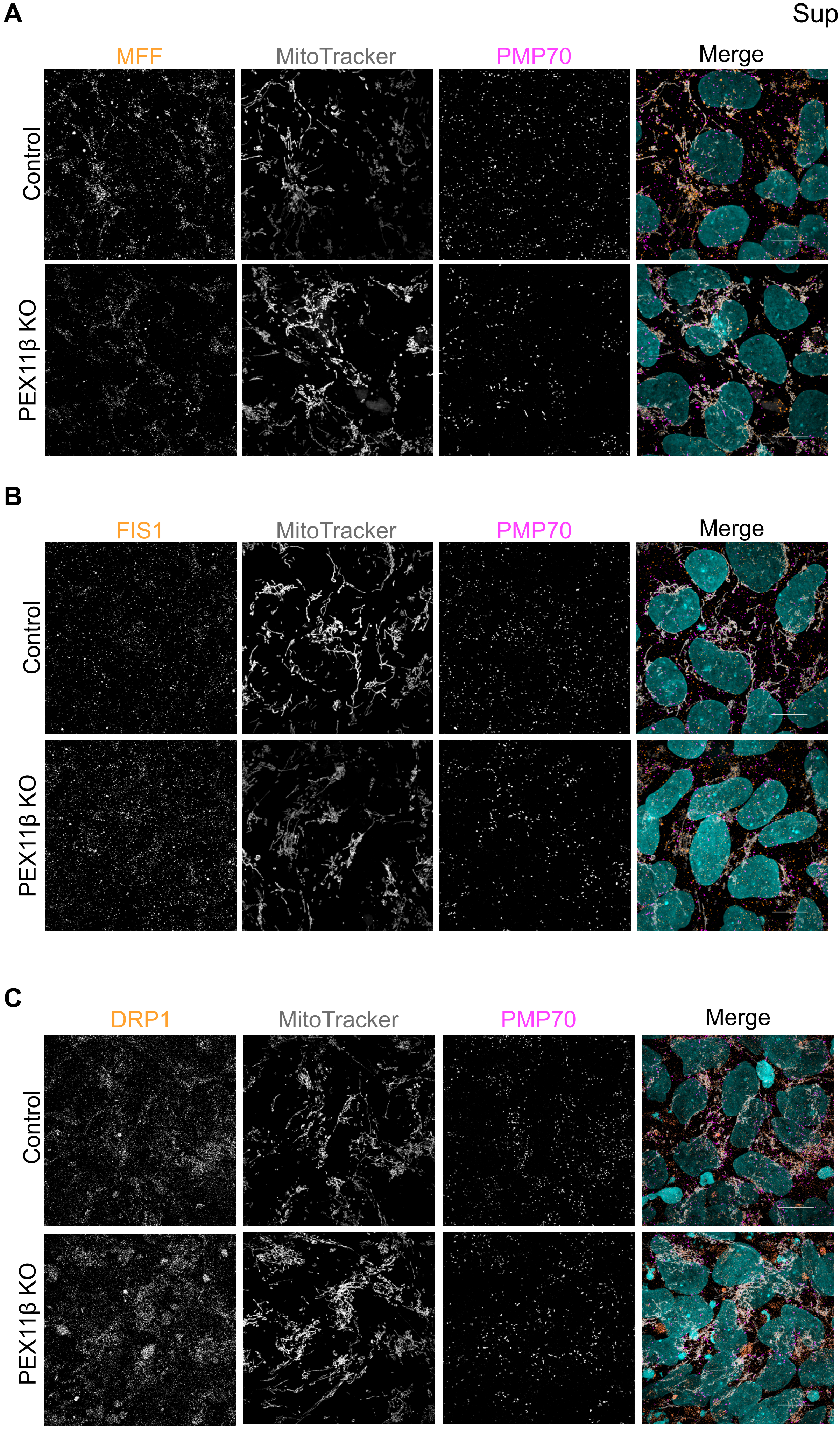

Supplementary Figure 7

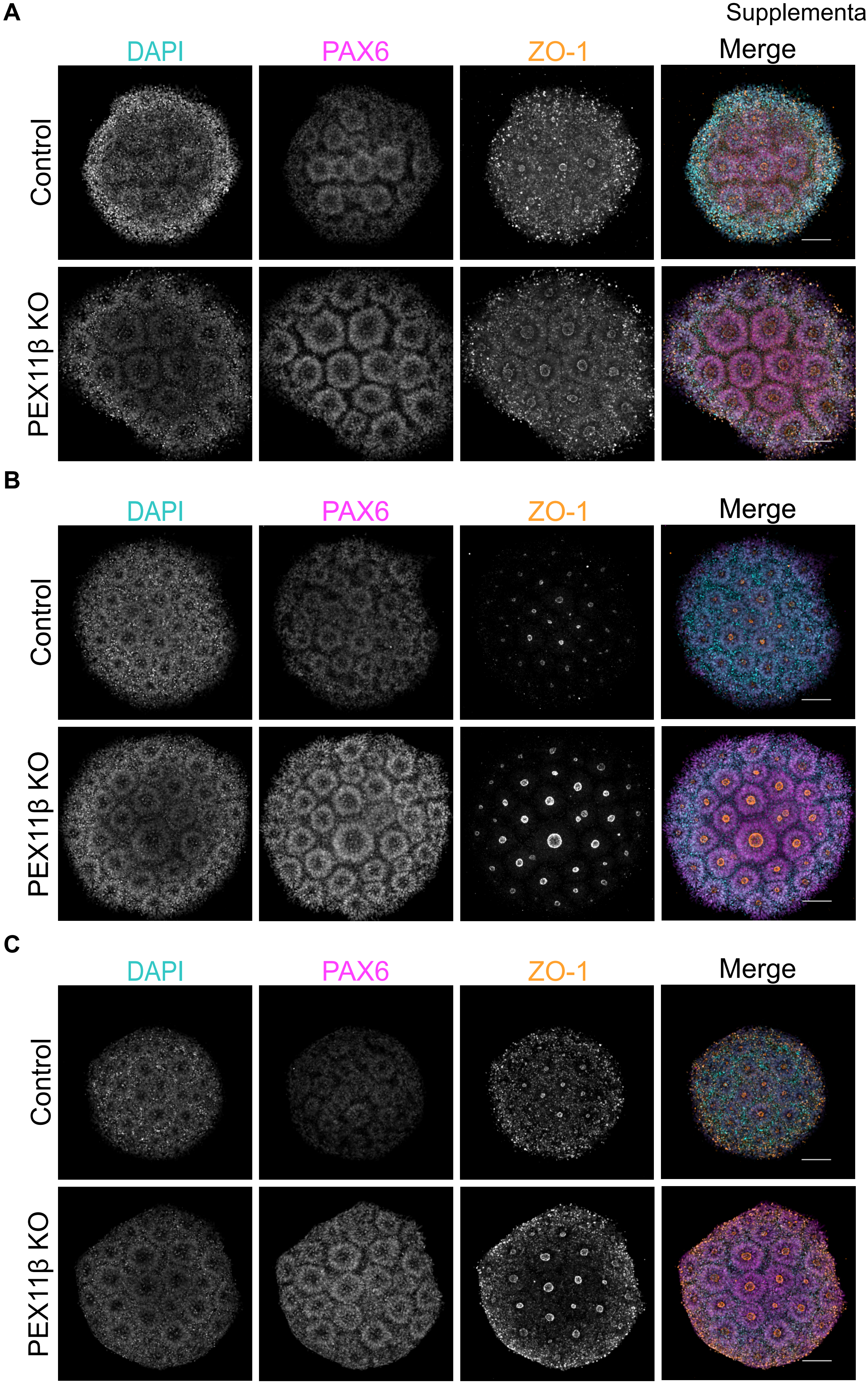
